# MAPK cascades mediating *Trichoderma brevicrassum* strain TC967 against phytopathogen *Rhizoctonia solani*

**DOI:** 10.1101/2021.09.01.458518

**Authors:** Yi Zhang, Wen-Ying Zhuang

## Abstract

*Trichoderma brevicrassum* strain TC967 is a novel biological control agent (BCA) against the plant pathogen *Rhizoctonia solani* and promotes plant growth. MAPK (mitogen-activated protein kinase) cascades were involved in biocontrol by *Trichoderma*, but functions of each MAPK in regulating biocontrol have not been characterized in one *Trichoderma*. In this study, we assembled and annotated the genome of strain TC967, and identified its three MAPK gene sequences. Functions of Fus3-, Slt2- and Hog1-MAPK in strain TC967 were dissected. The three MAPKs were all involved in hyphal growth. The Hog1-MAPK was essential for conidiation and tolerance to hyperosmotic stress. The Fus3- and Slt2-MAPK both mediated cell-wall integrity (CWI) and activities of chitinase and protease. The Fus3- and Hog1-MAPK mediated response to oxidative stress. Our biocontrol assays demonstrated that the Fus3- and Slt2-MAPK mutants were considerably more effective in disease control than the wild-type strain. RNA-seq analysis revealed that MAPK collectively played a major role in regulating biocontrol-related gene expressions, including of the genes in charge of secondary metabolism, fungal cell wall-degrading enzymes (FCWDEs) and small secreted cysteine-rich proteins (SSCPs).

**Author summary:** Soil-born fungal pathogens pose an emerging threat to crop production. *Trichoderma brevicrassum* strain TC967 has the ability to control the notorious phytopathogen *Rhizoctonia solani* and promote plant growth. In this study, we explored the functions of three-types of MAPK in mediating biocontrol process, and uncovered that Fus3-, Slt2- and Hog1-MAPK are involved in hyphal growth, conidiation, tolerance to hyperosmotic stress, cell-wall integrity, activities of chitinase and protease, and response to oxidative stress. Biocontrol ability of strain TC967 was accelerated after deletion of the Fus3- and Slt2-MAPK genes. MAPK collectively played a major role in regulating biocontrol-related gene expressions as revealed by RNA-seq analysis. To our knowledge, this is the first report of the functions of MAPKs in regulating biocontrol in one *Trichoderma*. Our results provide a reference for improvement of biocontrol ability of *Trichoderma* strains from the view of MAPK cascades.

## Introduction

*Trichoderma* strains are acted as very effective biological control agents (BCAs), and have been used for controlling a variety of phytopathogens [1, 2]. They possess different antagonistic strategies, competition, antibiosis, mycoparasitism, and induced resistance, for nutrients and niches as well as induction of enhancing systemic resistance and induced resistance responses in plants, which are suggested as the mechanisms of *Trichoderma* controlling pathogen infection process and disease development [3, 4]. Currently, 375 *Trichoderma* species are known in the world [5]. According to incomplete counting, about 25 *Trichoderma* species have the potential for effective control of diseases caused by more than 100 fungal plant pathogens, and most of them were commercialized as BCAs used for plant disease control [1, 2]. *Trichoderma brevicrassum* strain TC967 was recently isolated from soil of the Tibetan Plateau and able to enhance cucumber resistance against infection of the phytopathogen *Rhizoctonia solani*, and its efficiency was similar to the commercial BCA *T. harzianum* T22 [6, 7].

Mycoparasitism is one of the major mechanisms accounting for antagonistic activity of *Trichoderma* species [8]. Upon contacting with hyphae of phytopathogens, hydrolytic enzymes secreted by *Trichoderma*, such as chitinases, glucanases and proteases, were induced. These enzymes degraded cell walls of the pathogens following assimilation of their cellular contents [9, 10]. Furthermore, overexpression of the hydrolytic enzymes in *Trichoderma* resulted in more efficient biocontrol to inhibit phytopathogens [11–13]. Recent studies have shown that, except hydrolytic enzymes, secretomes of *Trichoderma* during interaction with host frequently contain a diversity of small secreted cysteine-rich proteins (SSCPs) [14–16]. The SSCPs might play a role in colonization, promotion plant growth and modulation of defense response [17, 18].

Multiple signal transduction pathways were proved participating in *Trichoderma* against phytopathogens, such as G-protein signaling, cAMP pathways and mitogen-activated protein kinase (MAPK) cascades [19–21]. MAPK cascades are highly conserved in fungi, plants and animals, and involve in transmission of extracellular and intracellular signals through regulating transcription factor by phosphorylation cascade [22, 23]. The signal transduction processes of MAPK cascades are started with sensing of environmental stimuli by receptors and proteins anchored to cell membrane [23]. The upstream elements of MAPK cascades, MAPK kinase kinase (MAPKKK), accept the signals, and which in turn activate MAPK kinase (MAPKK) by phosphorylation of serine/threonine residues. Subsequently, the MAPKs are activated by the phosphorylation, and finally give rise to the activation of transcription factors that induce or repress genes involved in cellular adaptation or response to the sensed stimuli [24, 25]. In fungi, the MAPK cascades have been well studied on *Saccharomyces cerevisiae* in which five types of MAPK were verified in relation to pheromone responses, filamentation growth, cell-wall integrity (CWI), osmotic stress, cell growth and conidiation [26, 27]. Accordingly, the genomes of *Trichoderma* species encode three types of MAPK. They belong to the so-called pathogenicity Fus3, cell-wall integrity Slt2, and osmoregulatory Hog1 [10, 28]. The involvement of MAPK in biocontrol of *Trichoderma* has attracted much attention. Deletion of Fus3-MAPK gene *tvk1* of *T. virens* enhanced biocontrol activity against *R. solani* and *Pythium ultimum* due to the production of more lytic enzymes [21]. In *T. atroviride*, the *tmk1* (Fus3-MAPK gene) deletion reduced mycoparasitic activity to *R. solani* and *Botrytis cinerea*, but the mutants displayed a higher inhibition ability against *R. solani* to protect bean plants [22]. However, deletion of Slt2-MAPK gene *tmkB* of *T. virens* caused cell-wall integrity defects and loss of antagonism against *Sclerotium rolfsii* but not *R. solani* and *Pythium* spp. [29]. Similar to Slt2-MAPK, Hog1-MAPK is also important for signaling in biocontrol, for example, deletion of Hog1-MAPK gene *hog1* of *T. harzianum* strongly reduced the antagonistic activity against the phytopathogens *Phoma betae* and *Colletotrichum acutatum* [29]. Therefore, Slt2- and Hog1-MAPK are paid less attention because the mutants of these genes displayed poor biocontrol ability.

In order to dissect the regulatory roles of the three types of MAPK of *T. brevicrassum* strain TC967 in biocontrol, we acquired the whole-genome of strain TC967; constructed the mutants of the three MAPKs; analyzed the characterizations of the mutants in growth, conidiation, stress tolerance and secondary metabolisms; and conducted an RNA-sequencing based global gene expression analysis of functions of the three MAPKs induced by *R. solani*. Such system and genome-wide characterization of the three MAPKs mediated biocontrol in a single *Trichoderma* strain has not been documented. Our data thus provides a complete insight into the function of MAPK-mediated biocontrol, and contributes to further improvement of biocontrol capacity of *Trichoderma* from the respect of MAPK signaling pathway.

## Results

### Features of strain TC967 genome assembly

The genome of *T. brevicrassum* strain TC967 was sequenced using the Illumina platform with 100× coverage, and the genomic sequence was assembled by Velvet. The genome of strain TC967 was 38.2 Mb in size across 105 scaffolds, with 49.57% GC content and maximum scaffold length of 3882954 bp (N50 length 1799946 bp) (Table 1). The whole-genome sequence of strain TC967 was deposited at GenBank under the accession JAEFBY010000000.

**Table 1.** Genome features of *Trichoderma brevicrassum* strain TC967.

| Genome feature | Value |
| --- | --- |
| Size (Mb) | 38.2 |
| Scaffolds | 105 |
| Scaffold N50 length (bp) | 1799946 |
| Scaffold maximum length (bp) | 3882954 |
| Scaffold minimum length (bp) | 1000 |
| GC content (%) | 49.57 |
| Genes | 11647 |
| rRNA genes | 44 |
| tRNA genes | 197 |
| Pseudogenes | 3 |

In the genome of strain TC967, a total of 11647 genes were predicted by using EVM combining with the *ab initio* and homology-based sequence annotation. The genome included 44 rRNA genes, 197 tRNA genes, and 3 pseudogenes (Table 1). The analysis of Nr annotation were executed, and the major proteins of *T. brevicrassum* strain TC967 mapped to other *Trichoderma* strains, such as 76.34%, 8.58% and 6.54% matched the proteins of *T. virens*, *T. reesei* and *T. atroviride*, respectively (Fig 1). The strains of *T. virens* and *T. atroviride* are commercially formulated biocontrol agents that are effectively employed against the soilborne phytopathogens *R. solani*, *S. rolfsii* and *Pythium* spp. [1, 30, 31].

**Fig. 1.**
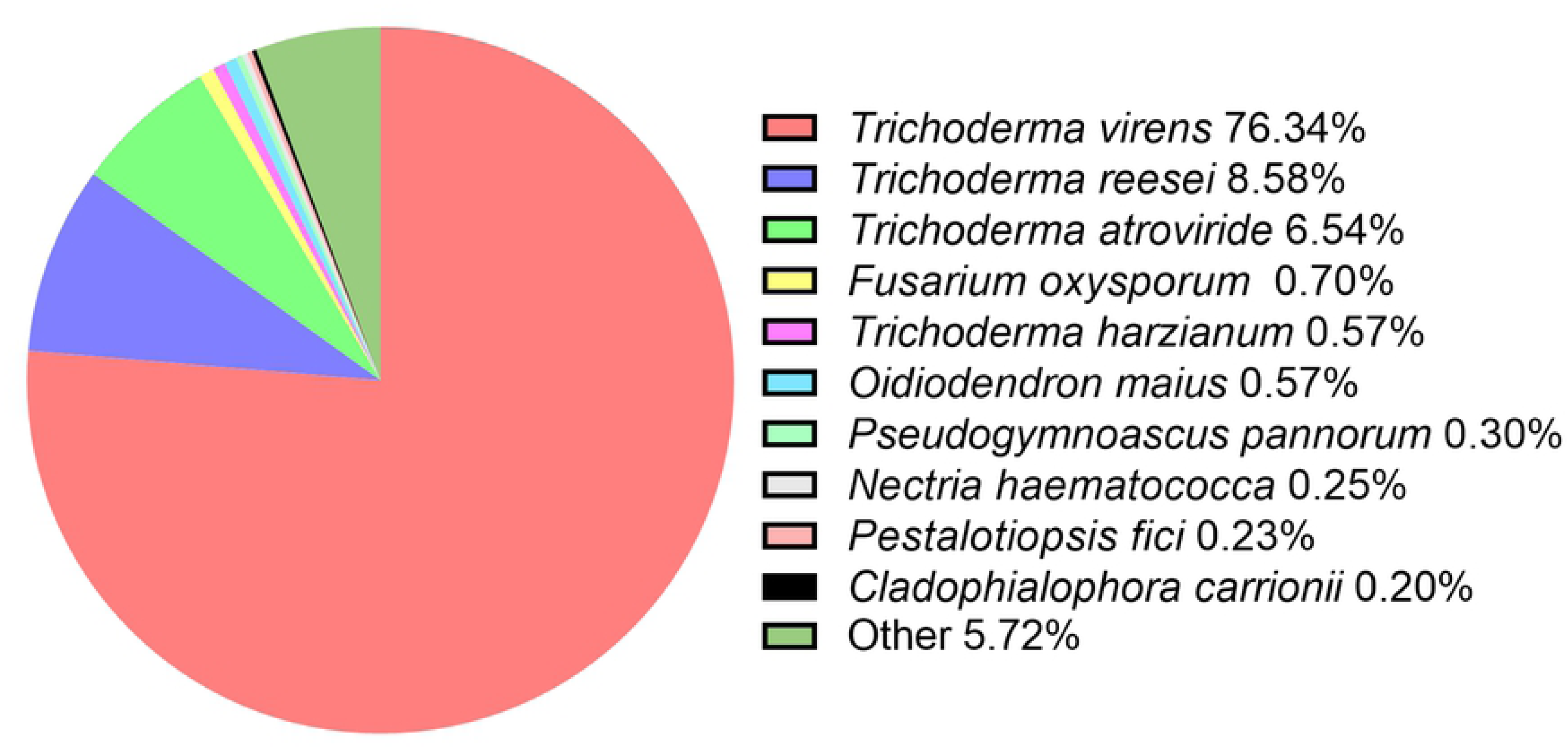
Non-redundant (Nr) classification analysis of the encoded proteins in the genome of *Trichoderma brevicrassum* strain TC967.

### Identification of *mapk* genes of strain TC967

Availability of the complete *T. brevicrassum* strain TC967 genomic sequences has made it possible for identifying the MAPK proteins in this strain. Based on the sequences of MAPK in *T. virens* Gv29-8, BLAST software was used to identify the *mapk* sequences. Three types of homogeneous MAPK genes were found in strain TC967, and named as *tbmk1*, *tbmk2* and *tbmk3*, respectively.

The MAPK is highly conserved across the fungal kingdom [23]. Similarly, the *mapk* sequences of *T. brevicrassum* strain TC967 had the same features as that of the investigated fungi. For example, there are similar number of exons and amino acids in each MAPK type (Table 2). Phylogenetic analysis of the predicted MAPKs of strain TC967 correlated with the previously characterized MAPK proteins indicated that TbMK1, TbMK2 and TbMK3 clustered with the Fus3, Slt2 and Hog1 types, respectively (Fig 2). Multiple alignment analysis of the predicted MAPK proteins showed that the three MAPKs of strain TC967 had conserved threonine and tyrosine residues in the TXY (Thr-X-Tyr) motif that is an activation loop, conformational change could result in the full activation of the MAPKs [32]. Moreover, MAPKs have a modified CD domain DXXDEP (Asp-X-X-Asp-Glu-Pro) in the C-terminal region (S1 Fig). The residues D (aspartate) and E (glutamate) play critical roles in interacting with the amino acids K (lysine) and R (arginine) in MAPKKs [33].

**Fig. 2.**
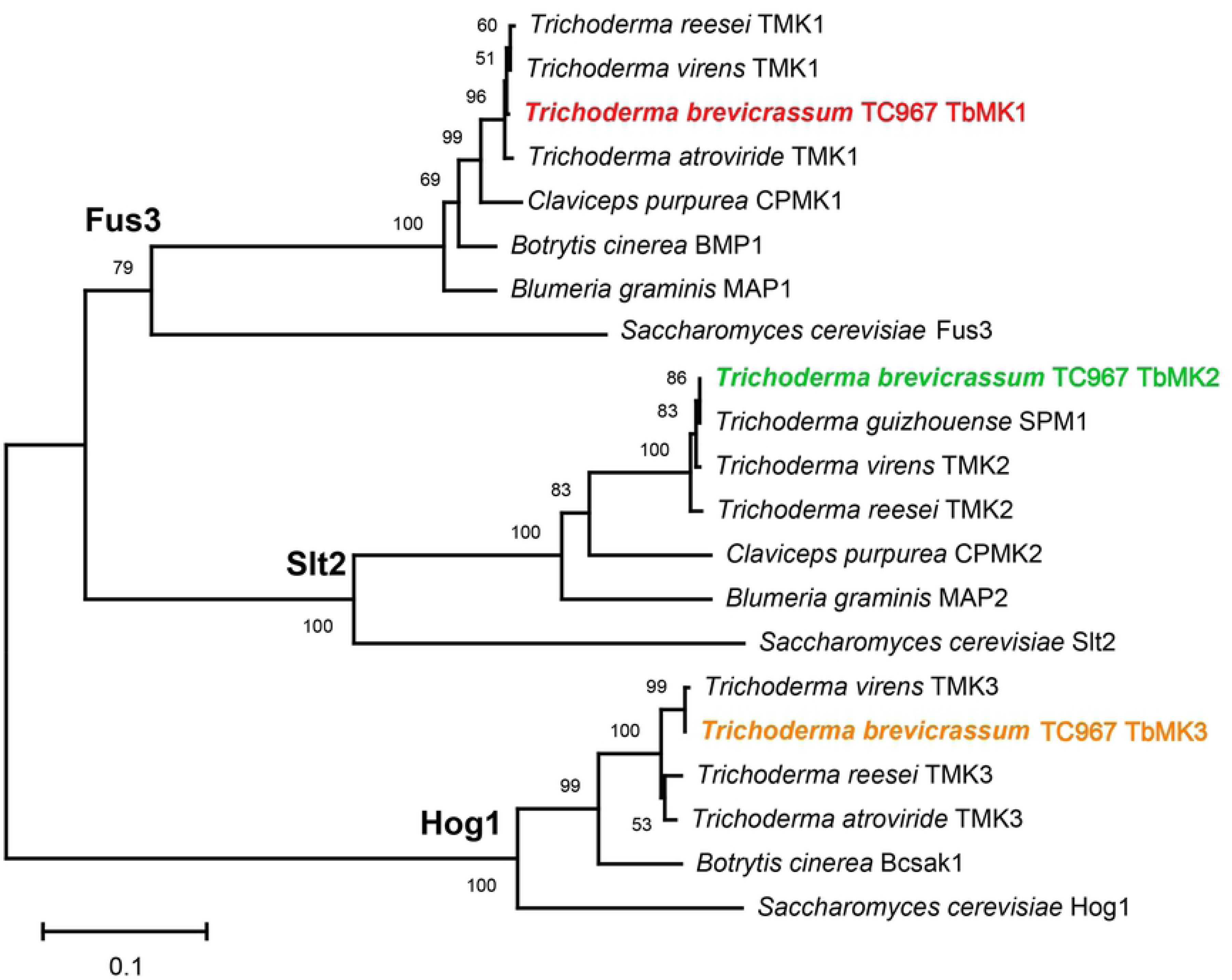
Phylogenetic relationships of MAPK protein sequences of fungal species. The phylogenetic tree was constructed by ML (maximum-likelihood) method using MEGA-X program. Numbers at the nodes indicate the ML bootstrap values with 1000 replicates. The red, green and orange represent Fus3-, Slt2- and Hog1-MAPK proteins in *Trichoderma brevicrassum* strain TC967, respectively. GenBank accession numbers were as follows: *Blumeria graminis* MAP1 (AAG53654) and MAP2 (AAG53655); *Botrytis cinerea* BMP1 (AAG23132) and BcSAK1 (AM236311); *Claviceps purpurea* CPMK1 (CAC47939) and CPMK2 (CAC87145); *Saccharomyces cerevisiae* Fus3 (AAA34613), Slt2 (CAA41954) and Hog1 (AAA34680); *T. atroviride* TMK1 (XP_013941981) and TMK3 (XP_013941593); *T. brevicrassum* strain TC967 TbMK1 (EVM0011518), TbMK2 (EVM0003677) and TbMK3 (EVM0007810); *T. guizhouense* SPM1 (OPB43978); *T. reesei* TMK1 (XP_006965066), TMK2 (XP_006969672) and TMK3 (XP_006962041); *T. virens* TMK1 (XP_013957486), TMK2 (XP_013959852) and TMK3 (EHK21342).

**Table 2.**
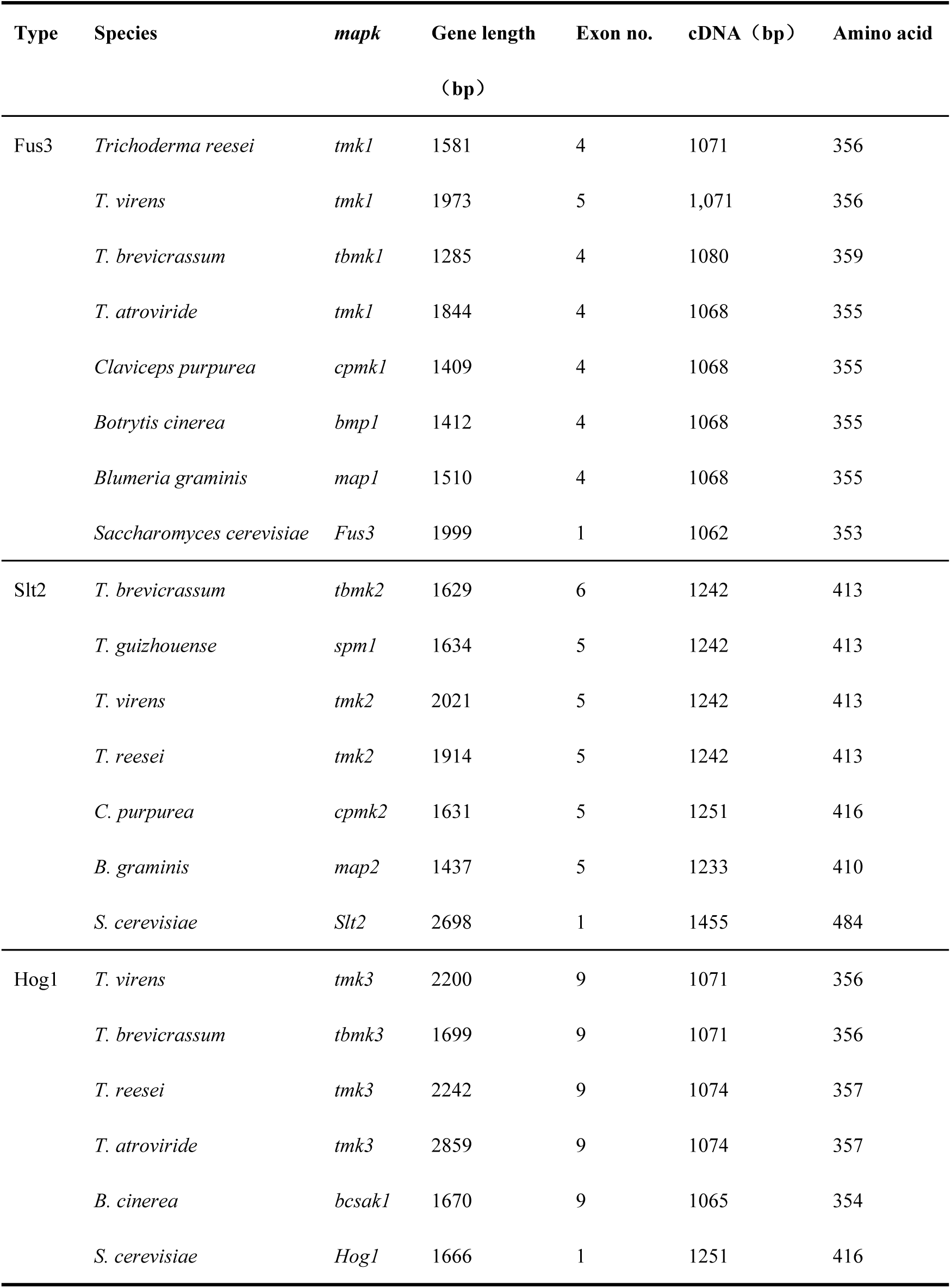
*mapk* sequence characteristics of some fungi.

| Type | Species | *mapk* | Gene length<br>(bp) | Exon no. | cDNA (bp) | Amino acid |
| --- | --- | --- | --- | --- | --- | --- |
| Fus3 | *Trichoderma reesei* | *tmk1* | 1581 | 4 | 1071 | 356 |
|  | *T. virens* | *tmk1* | 1973 | 5 | 1,071 | 356 |
|  | *T. brevicrassum* | *tbmk1* | 1285 | 4 | 1080 | 359 |
|  | *T. atroviride* | *tmk1* | 1844 | 4 | 1068 | 355 |
|  | *Claviceps purpurea* | *cpmk1* | 1409 | 4 | 1068 | 355 |
|  | *Botrytis cinerea* | *bmp1* | 1412 | 4 | 1068 | 355 |
|  | *Blumeria graminis* | *map1* | 1510 | 4 | 1068 | 355 |
|  | *Saccharomyces cerevisiae* | *Fus3* | 1999 | 1 | 1062 | 353 |
| Slt2 | *T. brevicrassum* | *tbmk2* | 1629 | 6 | 1242 | 413 |
|  | *T. guizhouense* | *spm1* | 1634 | 5 | 1242 | 413 |
|  | *T. virens* | *tmk2* | 2021 | 5 | 1242 | 413 |
|  | *T. reesei* | *tmk2* | 1914 | 5 | 1242 | 413 |
|  | *C. purpurea* | *cpmk2* | 1631 | 5 | 1251 | 416 |
|  | *B. graminis* | *map2* | 1437 | 5 | 1233 | 410 |
|  | *S. cerevisiae* | *Slt2* | 2698 | 1 | 1455 | 484 |
| Hog1 | *T. virens* | *tmk3* | 2200 | 9 | 1071 | 356 |
|  | *T. brevicrassum* | *tbmk3* | 1699 | 9 | 1071 | 356 |
|  | *T. reesei* | *tmk3* | 2242 | 9 | 1074 | 357 |
|  | *T. atroviride* | *tmk3* | 2859 | 9 | 1074 | 357 |
|  | *B. cinerea* | *bcsak1* | 1670 | 9 | 1065 | 354 |
|  | *S. cerevisiae* | *Hog1* | 1666 | 1 | 1251 | 416 |

In order to systematically study the characteristics of MAPK participation in biocontrol in strain TC967, the entire predicted ORF (open reading frame) of each *mapk* was deleted using a homologous recombination strategy. The details of the gene deleted and subsequent complementation are described in S2 Fig. Two mutants and one complemented mutant of each *mapk* were randomly chosen from all mutants using in subsequent test.

### Growth, hyphal morphology and conidiation of *mapk* mutants

The *mapk* deletion mutants were characterized for their defects in developments. All the *mapk* mutants showed significant slower growth rates compared with the wild and complemented mutant strains. However, the growth rate of Fus3-MAPK gene mutants (*Δtbmk1-6* and *Δtbmk1-8*) were faster than that of Slt2-MAPK gene (*Δtbmk2-4* and *Δtbmk2-5*) and Hog1-MAPK gene (*Δtbmk3-2* and *Δtbmk3-19*) mutants (Fig 3A and B). The hyphal morphology has changed after deletion of *mapk* genes. The mutants of Fus3-MAPK and Hog1-MAPK genes produced more branches at the hyphal tips with a normal width. The vegetative hyphae of Slt2-MAPK gene deletion mutants had thinner branches at the hyphal tips than that of the wild and complemented mutant strains (Fig 3). The results indicate that Slt2- and Hog1-MAPK are more critical for vegetative growth than Fus3-MAPK.

**Fig. 3.**
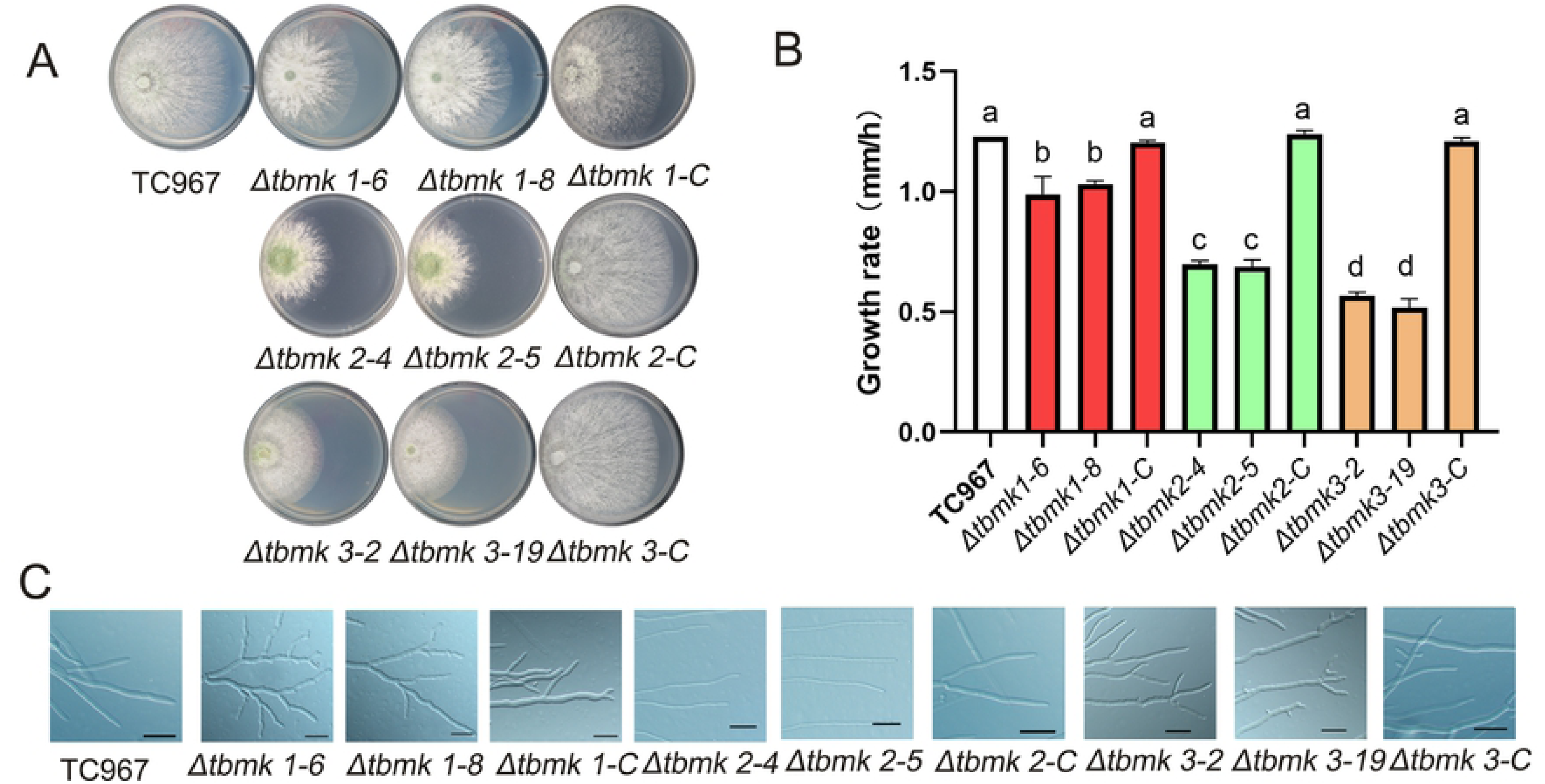
Growth rate and hyphal morphology of the wild and mutant strain of *Trichoderma brevicrassum* strain TC967. (A) Colony morphology of the wild and mutant strains of TC967 on PDA medium. *1-C*, *2-C* and *3-C* indicating the complemented strains. (B) Statistics of growth rate of the tested strains. The bars represent the averages of standard deviations of three biological replicates, different letters indicate statistically different (Fisher’s LSD, *p* < 0.05). (C) Hyphal morphology of the wild and mutant strains of TC967, bar = 50 μm.

To evaluate the involvement of *mapk* genes in conidiation, all the mutants and wild strain were cultured on PDA medium supplied with cycle light for 7 days. Our examinations showed that deletion of Hog1-MAPK gene led to seriously impaired conidiation. The Hog1-MAPK gene mutants (*Δtbmk3-2* and *Δtbmk3-19*) almost totally lost the ability of producing conidia (Fig 4A and C). The Fus3-MAPK gene (*Δtbmk1-6* and *Δtbmk1-8*) and the Slt2-MAPK gene (*Δtbmk2-4* and *Δtbmk2-5*) mutants did not significantly influence conidial production. Microscopic observations of the samples revealed that the conidiophores of Hog1-MAPK gene mutants were degenerated, which were much thinner and fewer, and conidia formed were rather sparse (Fig 4B). We also found that the Fus3- and Slt2-MAPK gene mutants lost concentric rings in colony, but the wild-type and complemented strains produced obvious concentric rings on PDA medium supplied with cycle light. In order to find out whether the *mapk* mutants are light dependence for conidiation, all strains were kept in a dark incubator. As the result, conidiation of Slt2-MAPK gene mutants (*Δtbmk2-4* and *Δtbmk2-5*) were sustained under dark condition (Fig 4D). In addition, Slt2-MAPK gene mutants produced massively conidia in submerged cultures (48 h) (Fig 4E), whereas no conidia were detected in cultures of the other mutants and wild strain even after 7 days. These facts suggest that Hog1-MAPK regulates conidiation, and Slt2-MAPK participates positive regulation of photoreception in the process of conidiation.

**Fig. 4.**
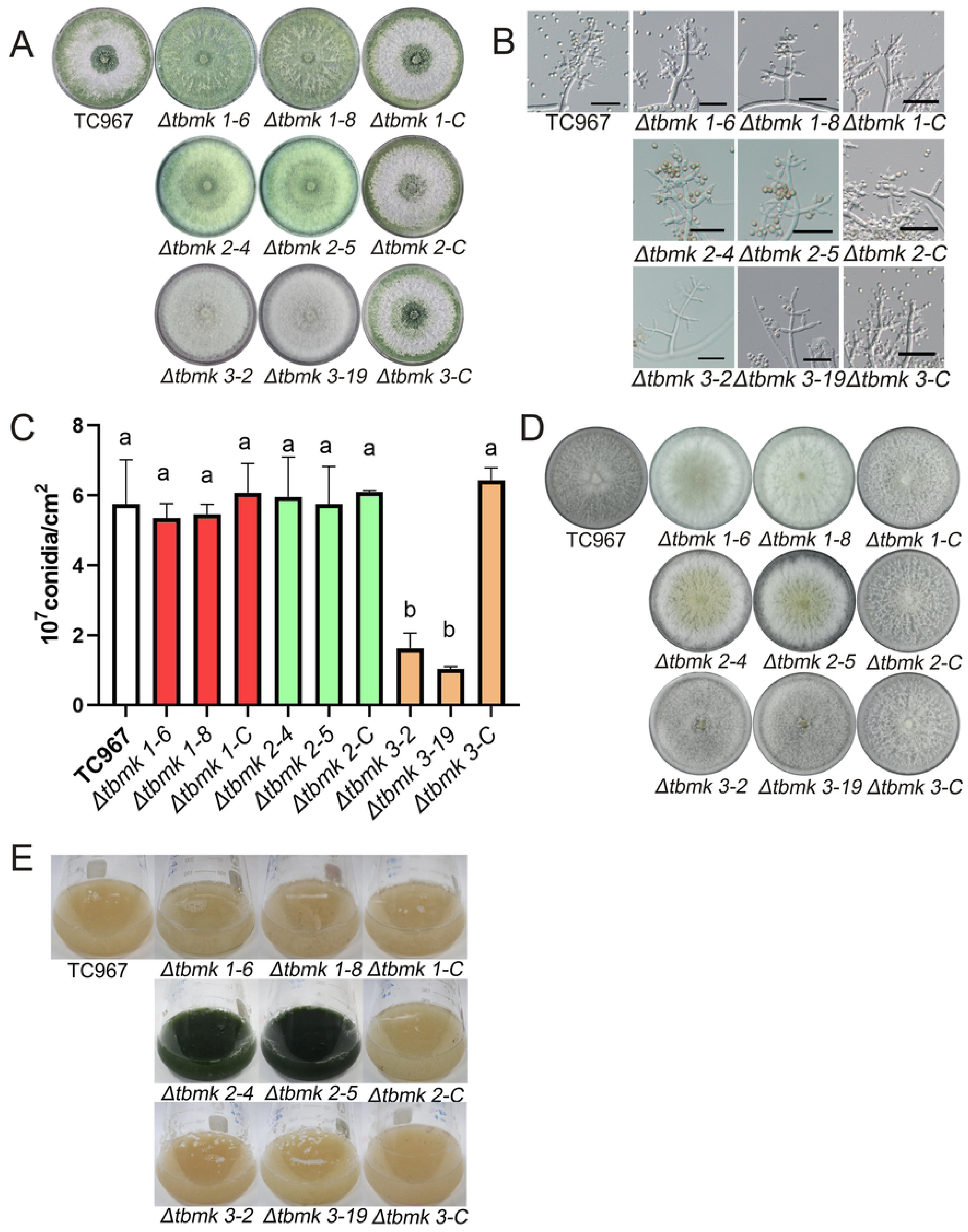
Conidiation of *mapk* mutant strains of *Trichoderma brevicrassum* stain TC967. (A) Colony and sporulation of the wild and mutant stains on PDA incubation at 25°C for 7 days under light/dark cycles. (B) Conidiophores of the wild and mutant strains, bars = 20 μm. (C) Number of conidia of each strain shown in (A). Bars represent the averages of standard deviations of three biological replicates, different letters on columns indicate statistically different (Fisher’s LSD, p < 0.05). (D) Colony morphology and conidiation of the wild and mutant strains on PDA incubation at 25°C for 7 days under dark condition. (E) Conidiation of the wild and mutant strains in PDB medium incubation at 25°C for 2 days under dark condition.

### Responses to abiotic stresses of *mapk* mutants

The cell-wall integrity of the wild and *mapk* mutant strains were investigated by testing the sensitivity to cell wall-damaging agent. The wild and *mapk* mutant strains were incubated on PDA medium containing Congo red (CR). The Fus3- and Slt2-MAPK gene mutants showed increased sensitivity to CR (Fig 5). The Hog1-MAPK gene mutants exhibited different degrees of tolerance to CR which was not harmful to growth of *Δtbmk3-2* and *Δtbmk3-19*. These results suggest that Fus3- and Slt2-MAPK genes were involved in maintaining the CWI.

**Fig. 5.**
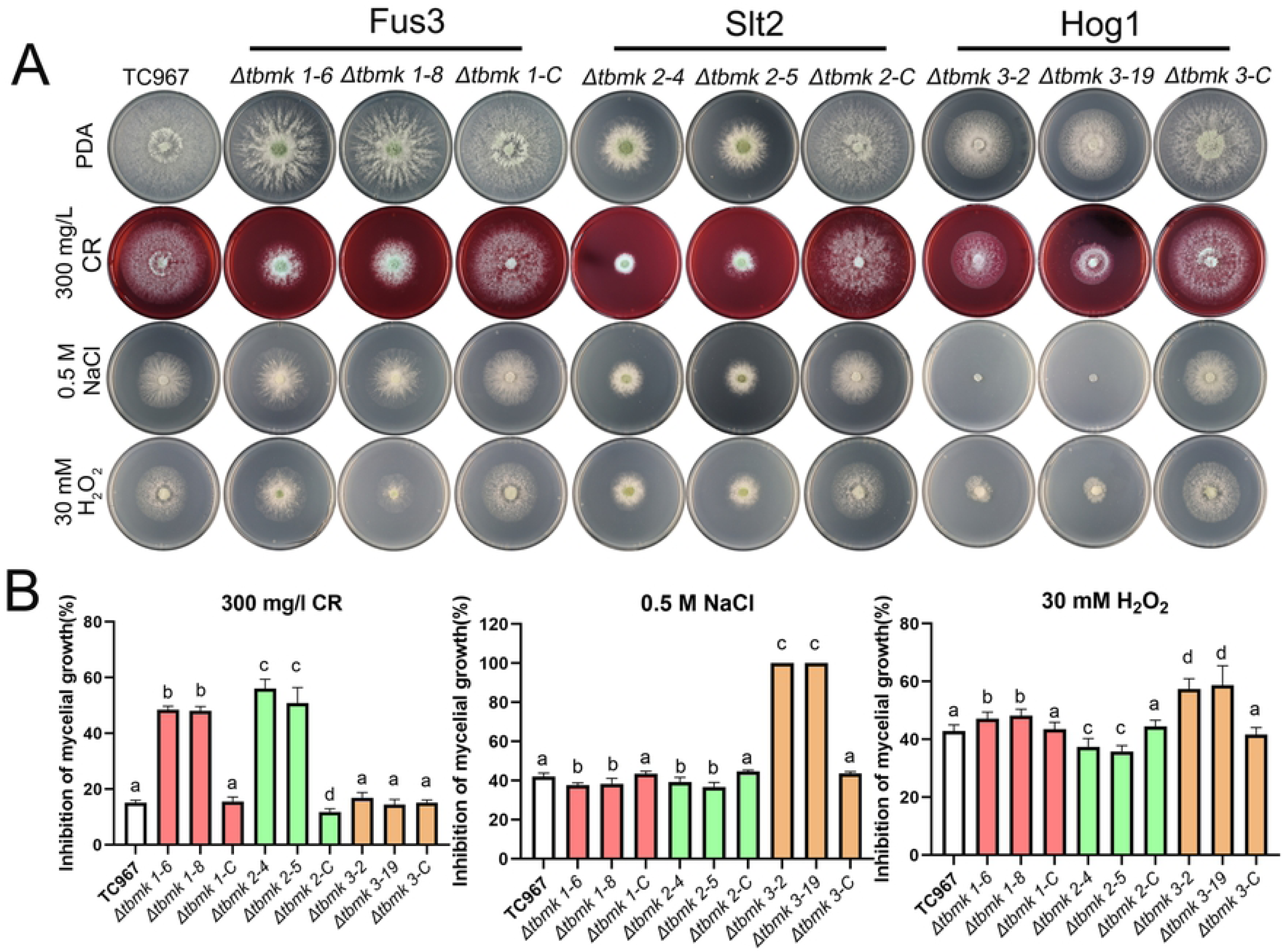
*mapk* mutant strains of *Trichoderma brevicrassum* strain TC967 response to abiotic stresses. (A) Sensitivity of Fus3-MAPK, Slt2-MAPK and Hog1-MAPK mutant strains to Congo red (CR), NaCl and H_2_O_2_ after incubation at 25°C for 3 days. (B) Mycelial growth inhibition of each strain showed in (A). White columns are the wild strain, red columns are Fus3-MAPK gene mutants, light green columns are Slt2-MAPK gene mutants, and orange columns are Hog1-MAPK gene mutants. Bars represent the averages of standard deviations of three biological replicates, different letters indicate significantly different (Fisher’s LSD, *p* < 0.05).

To determine whether these three types of *mapk* gene mutants have a defective response to osmotic stress, all the mutants and wild strain were cultured on PDA medium supplemented with 0.5 M NaCl. The growth of Hog1-MAPK gene mutants (*Δtbmk3-2* and *Δtbmk3-19*) were completely restricted (Fig 5). Our examination tells that Hog1-MAPK is involved in tolerance to hyperosmotic stress.

Under oxidative stress (PDA medium supplemented with 30 mM H_2_O_2_), growth of Fus3-MAPK gene (*Δtbmk1-6* and *Δtbmk1-8*) and Hog1-MAPK gene mutants (*Δtbmk3-2* and *Δtbmk3-19*) were significantly inhibited. These suggest that both Fus3-MAPK and Hog1-MAPK are involved in mediating response to oxidative stress in strain TC967.

### Influencing on growth of *R. solani* by *mapk* mutants

Wild strain TC967 possessed obviously the effects on restricting growth of *R. solani* in the previous study [7]. In order to investigate biocontrol potential of the *mapk* mutants, dual cultures were set up to determine their confrontation against *R. solani* (S3 Fig). The Fus3-MAPK gene deletion mutants (*Δtbmk1-6* and *Δtbmk1-8*) were similar to the wild and complemented mutants, which overgrew *R. solani* within 7 days. However, the Slt2- and Hog1-MAPK gene mutants exhibited reduced ability, especially the Hog1-MAPK gene mutants which almost completely lost the ability of overgrowing *R. solani*.

To further explore the functions of *mapk* mutants in production of antifungal metabolites. Firstly, we examined the secondary metabolites produced by all the mutants and wild strain (Fig 6A and B). *Rhizoctonia solani* was transferred to the PDA plates preincubated with *Trichoderma* strains. The growth of *R. solani* was obviously inhibited in the presence of metabolites produced by Slt2-MAPK gene (*Δtbmk2-4* and *Δtbmk2-5*) and Hog1-MAPK gene mutants (*Δtbmk3-2* and *Δtbmk3-19*).

**Fig. 6.**
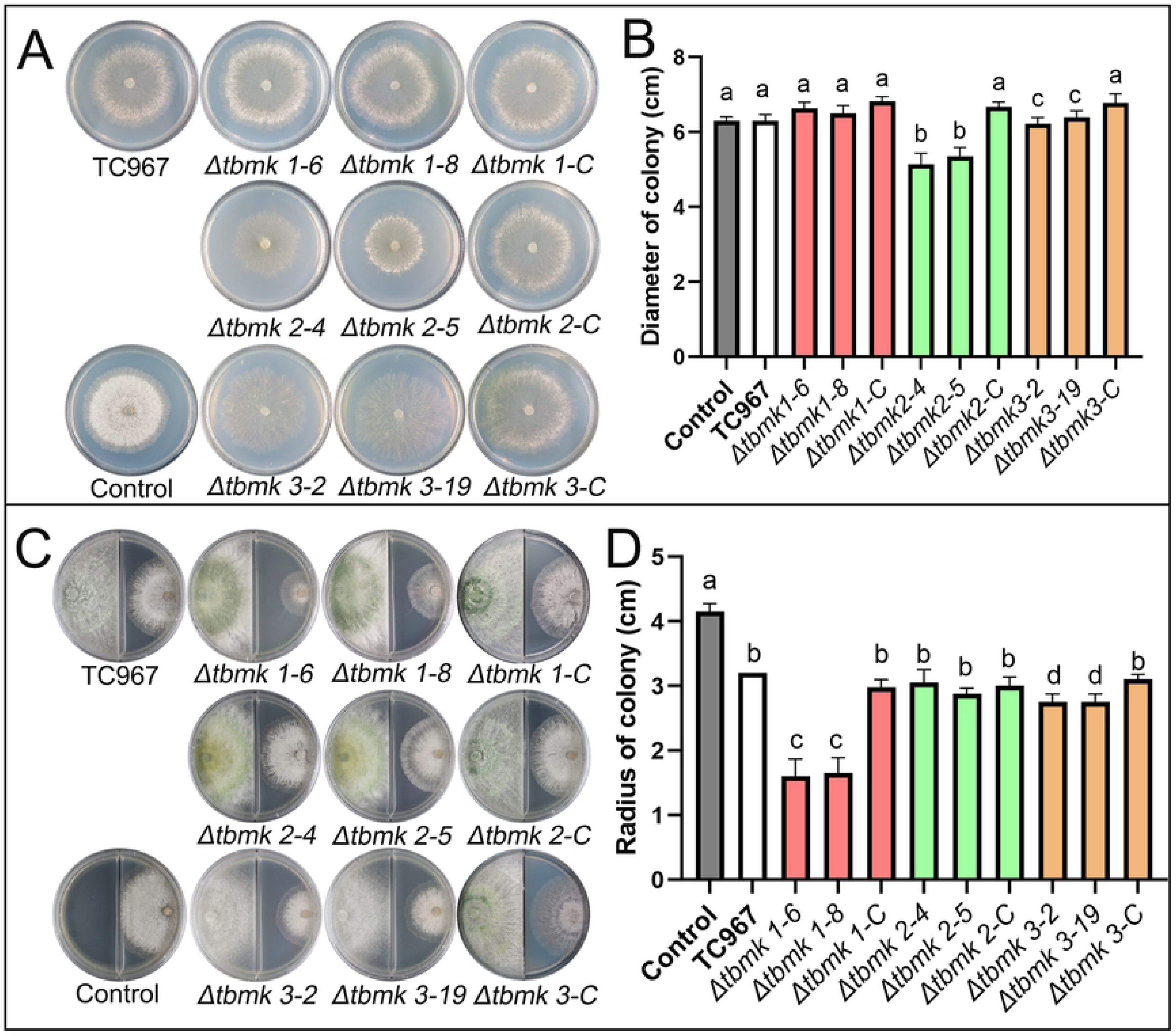
Inhibitory action of *mapk* mutants of *Trichoderma brevicrassum* strain TC967 antifungal metabolites on *Rhizoctonia solani*. (A) *R. solani* was grown on PDA plates containing secondary metabolites. Control, *R. solani* was cultivated on fresh PDA plates. All plates were incubated at 25°C for 2 days. (B) and (D) Mycelial growth inhibition of each strain shown in (A) and (C) respectively, bars represent the averages of standard deviations with three biological replicates, different letters are significantly different (Fisher’s LSD, *p* < 0.05). (C) Mycelial growth inhibition of *R. solani* exposed to VOCs produced by *Trichoderma* strains. *R. solani* (right) and *Trichoderma* strains (left) grown on PDA in I-plates at 25°C for 2 days. Control, only *R. solani* inoculated on one side of I-plates.

Volatile organic compounds (VOCs) are important antifungal metabolites. Growth of *R. solani* in the presence of VOCs from *Trichoderma* strains were tested. The control lacking exposure to *Trichoderma* VOCs grew an average of 4.15 cm after 2 days. Mycelial growth of *R. solani* in the presence of VOCs was all suppressed. Exposure to the suite of the VOCs emitted from the wild strain TC967 and Hog1-MAPK gene mutants (*Δtbmk3-2* and *Δtbmk3-19*) yielded 22.9% and 34.9% growth inhibition, respectively. The greatest inhibition was associated with exposure to VOCs emitted from Fus3-MAPK gene mutants (*Δtbmk1-6* and *Δtbmk1-8*), which inhibited the mycelial growth of *R. solani* by up to 61.4%. The inhibition of VOCs by the Slt2-MAPK gene mutants (*Δtbmk2-4* and *Δtbmk2-5*) appeared no significant difference from the wild strain and complemented mutants (Fig 6C and D). These results imply that Fus3-MAPK may negatively regulate VOCs produced by strain TC967, and deletion of this gene enhances growth inhibition of *R. solani* by the VOCs.

### FCWDEs activity of *mapk* mutants

Fungal cell wall-degrading enzymes (FCWDEs) were important secondary metabolites, which took part in hydrolyzing fungal cell walls [10]. Three types of FCWDE activities were tested, and the results were shown in Fig 7 and S4 Fig. The activities of chitinase and protease of Fus3-MAPK gene (*Δtbmk1-6* and *Δtbmk1-8*) and Slt2-MAPK gene mutants (*Δtbmk2-4* and *Δtbmk2-5*) significantly increased comparing with the wild strain and complemented mutants. As to the activity of β-1,3-glucanase, obvious difference was not detected between the mutants and wild strain (S4C Fig). From the above mentioned, we tentatively conclude that Fus3- and Slt2-MAPK negatively regulate the expressions of chitinase and protease-related genes.

**Fig. 7.**
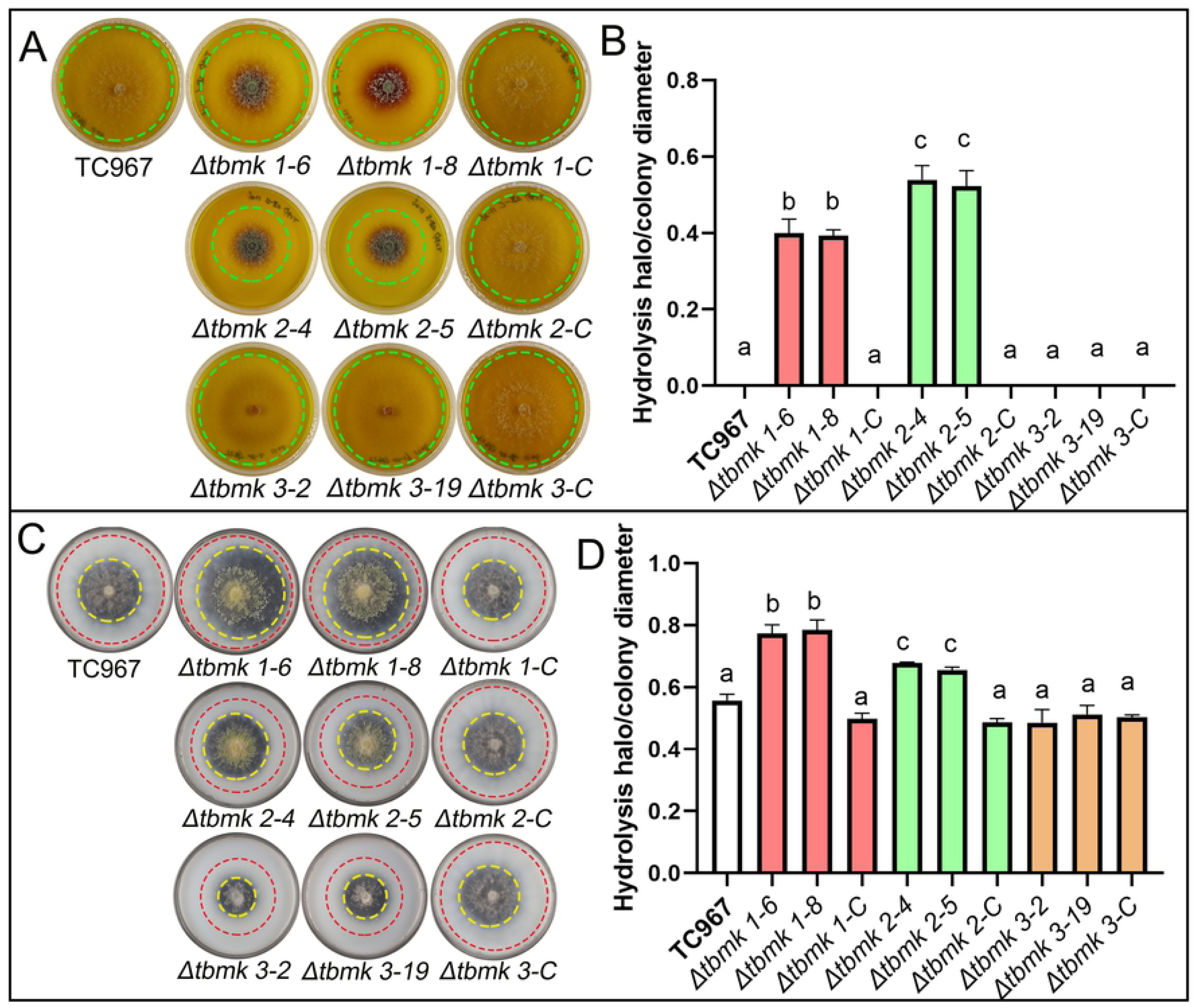
Chitinase and protease activities of *mapk* mutants of *Trichoderma brevicrassum* strain TC967. (A) Measurements of chitinase activity in Petri dishes containing the testing substrates, green dotted line indicating the edge of colony, purple area indicating the chitinase hydrolytic circle. (B) and (D) Ratio of hydrolysis halo to colony diameter shown in (A) and (C), respectively. Bars represent the averages of standard deviations of three biological replicates, different letters are significantly different (Fisher’s LSD, *p* < 0.05). (C) Measurements of protease activity in Petri dishes containing the testing substrates, red dotted line indicating the edge of colony, yellow dotted line indicating the protease hydrolytic circle.

### Biocontrol activity of *mapk* mutants

Biocontrol abilities of *mapk* mutants were tested *in vivo* against *R. solani*. Although protection of cucumber from infection of *R. solani* by wild strain TC967 appeared to have a good effect [Disease index (DI) 38.9%], the cucumber seedlings had lower morbidity when treated with the Fus3-MAPK gene mutant (*Δtbmk1*) (DI 33.33%) and Slt2-MAPK gene mutant (*Δtbmk2*) (DI 34.72%). The DI of the Hog1-MAPK gene mutant (*Δtbmk3*) (DI 41.67%) was less than the RS treatment (DI 59.72%), but it was higher than the Fus3- and Slt2-MAPK gene mutants. Overall, these results support that the Fus3- and Slt2-MAPK gene mutants enhanced cucumber plants to defend against *R. solani*.

### Overview of RNA-seq analysis of MAPK-mediated gene regulation induced by *R. solani*

After the *Trichoderma* strains recognizing *R. solani*, hyphae of *Trichoderma* coiled that of *R. solani*. During mycoparasitism, expressions of biocontrol-related genes of *Trichoderma* were induced by the pathogen. In order to explore the function of MAPKs in biocontrol, transcriptomes of the wild strain, Fus3-MAPK gene mutant (*Δtbmk1*), Slt2-MAPK gene mutant (*Δtbmk2*) and Hog1-MAPK gene mutant (*Δtbmk3*) were profiled by Illumina NovaSeq 6000 RNA-seq after hyphae of *Trichoderma* and that of *R. solani* contacted. A heatmap showed that high consistency between three biological replicates (R^2^ > 0.85) (S5 Fig). These draw the attention to the fact that disruption of any one of the three types MAPK genes does not affect expression level of a significant number of genes associated with biocontrol.

More than 93.64% of the resulted sequences in RNA library of the three types of MAPK gene mutants after contacting with *R. solani* (AC treatments) was mapped on genome of strain TC967 (S1 Table). The correlation between the RNA-seq and real-time qPCR results was highly significant (R^2^ >0.86) (S6 Fig), which confirms the reliability of RNA-seq results. Therefore, the resulted sequencing data is suitable for running further analysis.

The number of differentially expressed genes (DEGs) between *mapk* mutants AC and TC967 AC treatments was 1841 in *Δtbmk1*, 1686 in *Δtbmk2*, and 2123 in *Δtbmk3*. The “upregulated genes” are genes with higher expression levels in mutant AC than TC967 AC treatments, and the “downregulated genes” are that with lower expression levels. In *Δtbmk2* and *Δtbmk3* strains, numbers of the upregulated genes were approximately equal to that of downregulated genes. But, the number of upregulated genes was more than the downregulated ones in *Δtbmk1* strain (Fig 9A). There were more common downregulated genes (310) than the common upregulated ones (132) in the three MAPK mutants (Fig 9B). Based on the above information and combining with the results from pot experiment of *Δtbmk1* and *Δtbmk2*, these two strains have strong biocontrol ability (Fig 8), function of the upregulated DEGs of each treatment was examined in order to find the gene enrichment of the *mapk* mutants induced by *R. solani*. In *Δtbmk1* strain, DEGs were enriched in 14 GO (Gene Ontology) terms, including eight in molecular function (Fig 9C), four in biological process (Fig 9D), and two in cellular component (Fig 9E) (FDR < 0.05). Seven of the 14 GO terms were related to hydrolase (GO:0016491, GO:0008171, GO:0016810, GO:0004497, GO:0016832, GO:0055114 and GO:0005975). For *Δtbmk2* strain, DEGs were associated with 19 GO terms, incorporating 12 in molecular function, three in biological process, and four in cellular component (FDR < 0.05). Except some of the above-mentioned hydrolases, DEGs were also enriched in other hydrolases terms (GO:0016491, GO:0008171, GO:0016832, GO:0004365 and GO:0033897).

**Fig. 8.**
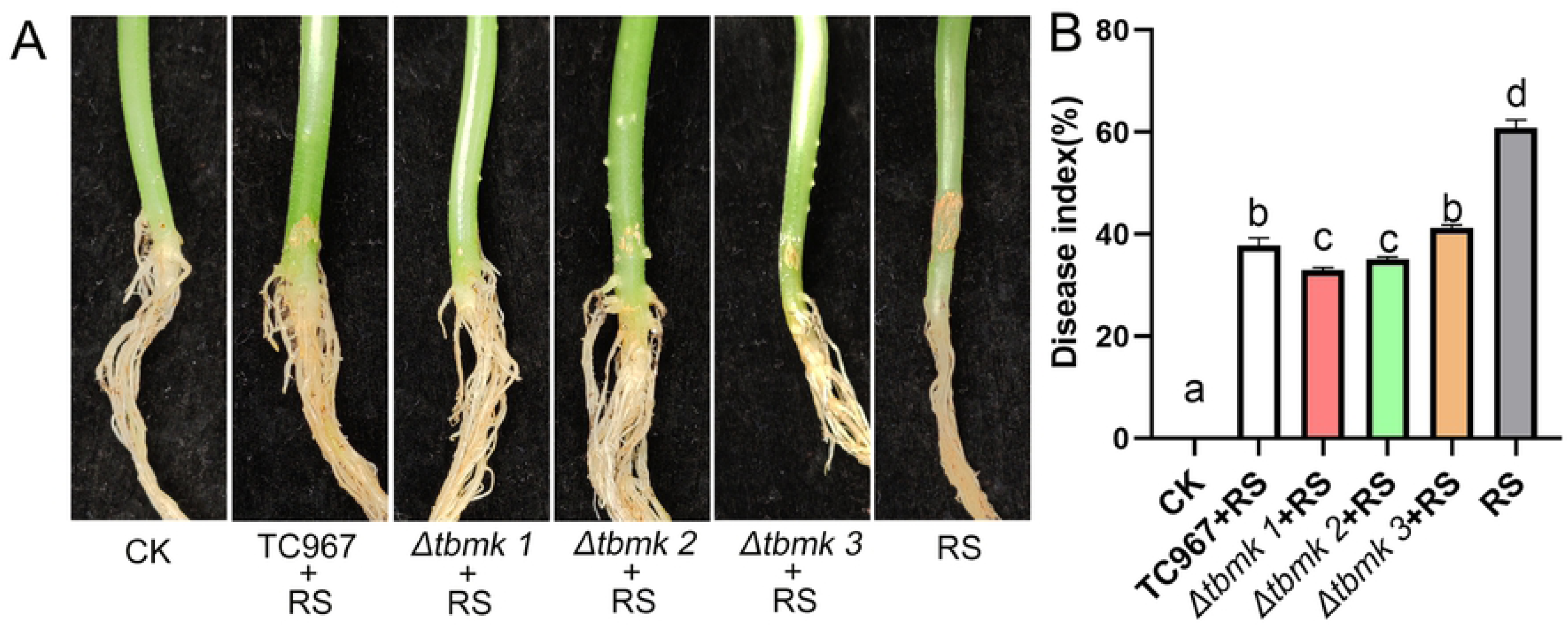
Control of *Rhizoctonia solani* infection on cucumber seedlings by *mapk* mutants of *Trichoderma brevicrassum* strain TC967. (A) Disease symptom of cucumber seedlings caused by *R. solani*. CK, not treated with any fungi; RS, treated with *R. solani* only; *Trichoderma* strains + RS, pretreated with *Trichoderma* strains then inoculated with *R. solani*. (B) Disease index of cucumber seedlings.

**Fig. 9.**
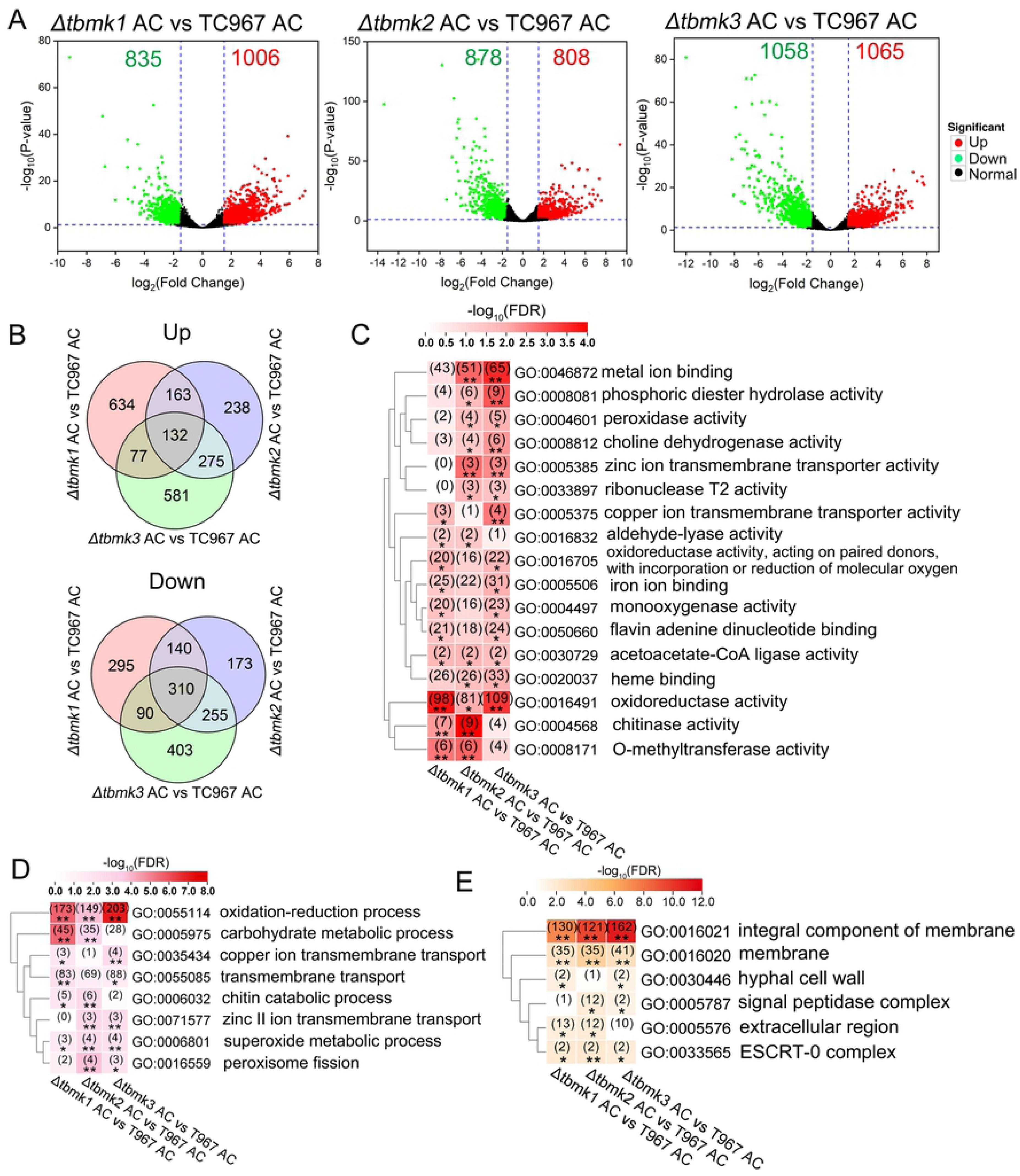
Statistical analysis of DEGs in *mapk* mutants of *Trichoderma brevicrassum* strain TC967 induced by *Rhizoctonia solani*. (A) Volcano plot of DEGs in three types *mapk* mutants after contacting with hyphae of *R. solani* (*Δtbmk1* AC, *Δtbmk2* AC and *Δtbmk3* AC), as compared with the wild strain of TC967 after contacting with *R. solani* (TC967 AC). (B) Venn diagrams show overlap of the upregulated genes (Up) and downregulated genes (Down) of *mapk* mutants of TC967 induced by *R. solani*. GO items in molecular function (C), biological process (D) and cellular component (E) were enriched among upregulated DEGs between the mutants and wild strain of TC967. Number in parenthesis indicates the number of genes in that category of each treatment. **FDR < 0.01, *FDR < 0.05.

### MAPK-mediated regulation of expression of genes related to biocontrol

To determine the three MAPK signal transduction pathways involved in biocontrol process, we analyzed gene expressions of secondary metabolism and FCWDEs. The genome of strain TC967 encoded 11 non-ribosomal peptide synthase (NRPS) clusters and NRPS-like clusters (S7 Fig and S2 Table). Eight of the nine NRPS and NRPS-like cluster contained basic AMP-binding domain except for NRPS-like 4, which only carried peptidyl-carrier protein domain and thioesterase domain (S7 Fig). Eight T1PKS (type I polyketide synthase)-NRPS and 16 T1PKS clusters were identified in TC967 genome, they all had typical beta-ketoacyl-synthase domain (S7 Fig and S2 Table). Six terpene clusters were also found in the genome (S2 Table). After hyphae of *Trichoderma* strain contacting with that of *R. solani*, the expressions of core genes of the clusters were identified among the three *mapk* mutants in comparison with the wild strain. In *Δtbmk1* strain, 26 genes were upregulated, and six were downregulated. The numbers of upregulated and downregulated genes in *Δtbmk2* strain were the same as that in *Δtbmk1* strain. However, most of the core genes of T1PKS clusters were downregulated in *Δtbmk3* strain (Fig 10A). It means that deletion of *tbmk1* and *tbmk2* enhanced the gene expressions of secondary metabolism. And it coincides with the results of the *Δtbmk1* and *Δtbmk2* strains which have higher antifungal activities of secondary metabolites than that in the *Δtbmk1* and wild strains.

**Fig. 10.**
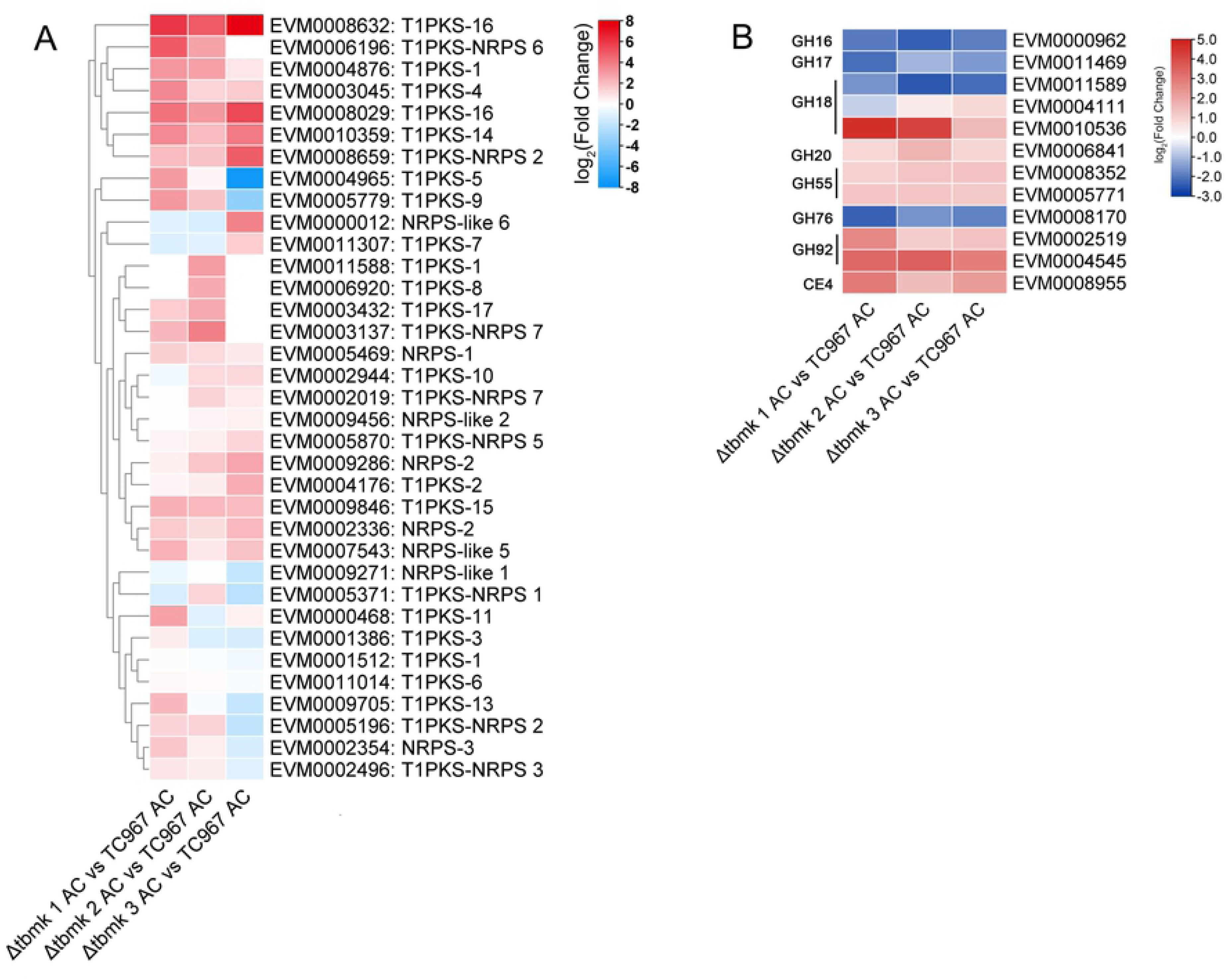
Expression of core genes related to secondary metabolism and FCWDEs in *mapk* mutants of *Trichoderma brevicrassum* strain TC967 induced by *Rhizoctonia solani*. (A) Heatmap of DEGs in *mapk* mutants induced by *R. solani*. (B) Expression of FCWDEs in CAZymes detected after *Trichoderma* strains contacting with *R. solani*.

*Trichoderma brevicrassum* strain TC967 encoded 88 FCWDEs distributed in 21 families of CAZymes. Of these, 81 FCWDEs belonged to 13 glycoside hydrolase (GH) families, four in carbohydrate esterase (CE) families contained four FCWDEs, and three FCWDEs distributed in two cellulose-binding module (CBM) families (S3 Table). Analysis of the DEGs of all FCWDEs in TC967, 12 genes were significantly changed in the three *mapk* mutants compared with wild TC967 (FDR < 0.05). Seven genes were upregulated in the three *mapk* mutants, including of one gene in chitinase family (GH18), two in β-1,3-glucanase family (GH55), two in α-mannosidase family (GH92), and one in acetyl xylan esterase family (CE4). Meanwhile, four genes were downregulated in all the mutants, and respectively distributed in cell wall glucanase (GH16), GH18, and fructose-bisphosphate aldolase (GH76) (Fig 10B). These results reveal that the three *mapk* genes in TC967 have equal contribution to regulating gene expressions of FCWDEs.

### SSCPs enriched in the *mapk* mutants

Recent studies found that small secreted cysteine-rich proteins are unique for fungi. SSCPs have been investigated in fungi that interact with hosts and pathogens [18, 34]. The amino acids with small size (less than 300) and more than 4% content of cysteine residues were chosen for analysis. A total of 261 SSCPs were predicted in strain TC967, which account for 7.7% of the total predicted proteins. The GO enrichment analysis revealed that 159 SSCPs were enriched in molecular function, cellular component, and biological process (FDR < 0.05). And they were mainly distributed in carbon-sulfur lyase activity (GO: 0016846), metabolic process (GO: 0008152), and extracellular region (GO: 0005576) (Fig 11A and S4 Table).

**Fig. 11.**
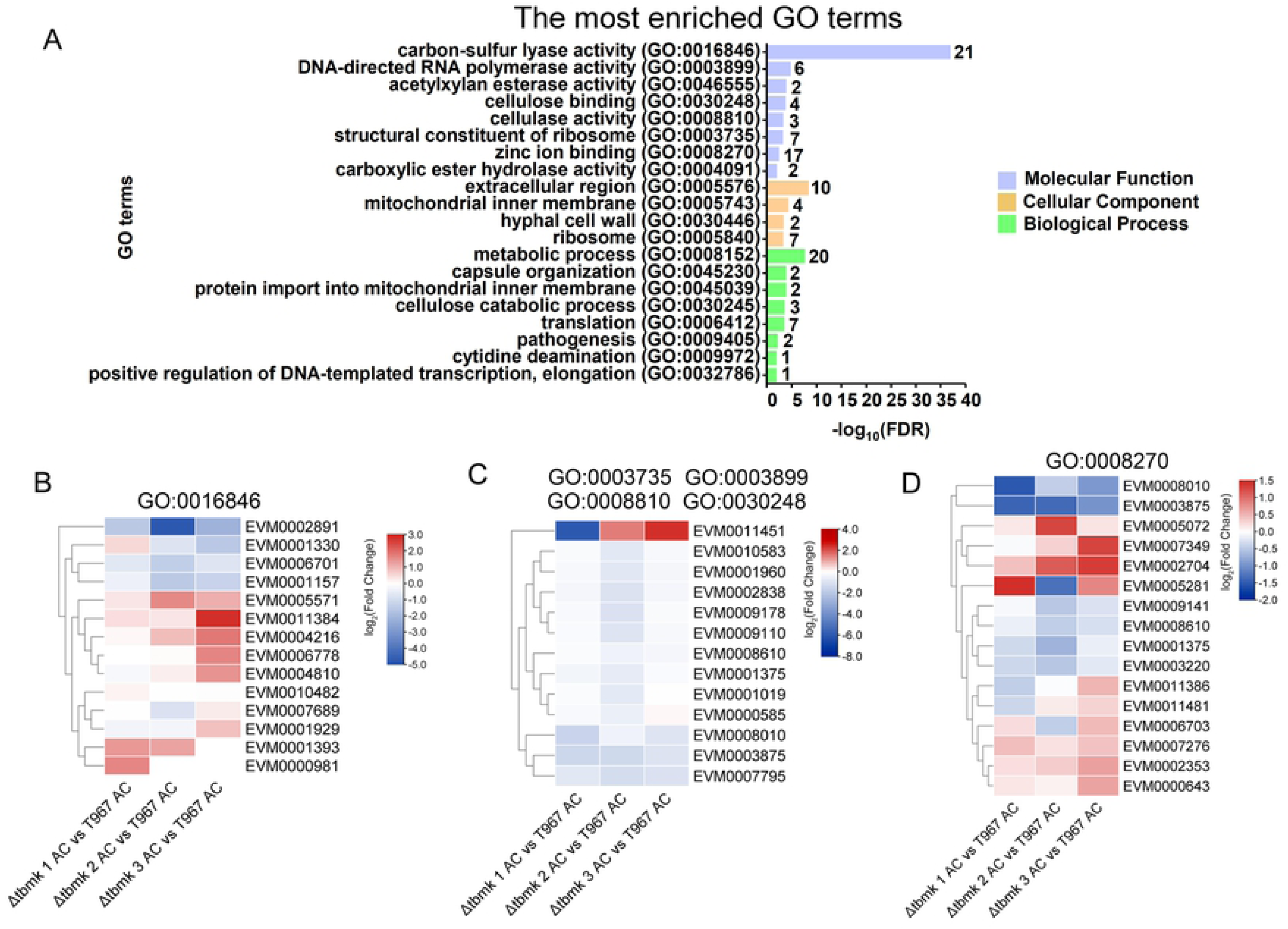
Small secreted proteins in the *mapk* mutants of *Trichoderma brevicrassum* strain TC967. (A) Enriched GO terms of small secreted proteins, the numbers following the columns indicate gene numbers of each GO term. (B)‒(D) DEGs analysis of genes enriched in molecular function. (B) Expression of the DEGs involved in GO:0016846. (C) Expression of the DEGs involved in GO:0003735, GO:0003899, GO:0008810, and GO:0030248. (D) Expression of the DEGs involved in GO:0008270.

Most of the SSCP genes involved in the molecular function term were upregulated in *Δtbmk3* strain (Fig 11B, C, D). However, only several putative glutathione-dependent formaldehyde-activating enzyme genes (EVM0001330, EVM0010482, EVM0001393 and EVM0000981) were upregulated in the other two mutant strains (Fig 11B, C, D). These results may be explained as *tbmk1* and *tbmk2* are involved in positive regulation of expression of most SSCPs genes, while *tbmk3* plays a negative regulatory role in most of the SSCP genes.

## Discussion

*Trichoderma brevicrassum* strain TC967 possessed prominent biocontrol ability against the phytopathogen *R. solani* in greenhouse pot experiment [7]. In the present work, we sequenced the whole genome of strain TC967, the size of its genome was approximately 38.2 Mb, which is similar to that of the other sequenced *Trichoderma* strains. Three different types of *mapk* genes were found, which are corresponding to the characterizations of verified MAPKs.

MAPK cascades play an important role in cellular regulation of fungi, but the functions of three MAPKs mediating biocontrol in one *Trichoderma* have not been characterized. Using reverse genetic methods, the roles of MAPKs in maintaining fungal characteristics and mediating genes expressions in biocontrol strain of *Trichoderma* were systematically explored. The Fus3-MAPK was an essential protein for filamentous growth, pheromone response, and mating of fungi [23, 35, 36]. Deletion of Fus3-MAPK gene (*tbmk1*) slightly effects the radial growth of strain TC967, and has no obvious effect on production of conidia. The maintained CWI was broken after deletion of *tbmk1*, but its antifungal activities by producing VOCs and FCWEDs were obviously improved comparing with the wild strain. Similarly, the Fus3-MAPK gene mutants showed higher activity of FCWEDs in other *Trichoderma* strains, such as *T. virens* Gv29-8 and *T. atroviride* P1 [21, 22]. Unlike the Fus3-MAPK gene mutant (*Δtvk1*) of *T. virens* Gv29-8, which produced massively conidia in submerged cultures [21, 37], the conidia produced by Fus3-MAPK deficient strain TC967 is similar to the wild type. Unexpectedly, the Slt2-MAPK gene deletion mutant of strain TC967 still produced massive conidia even in submerged cultures at dark condition, which is similar to the Slt2-MAPK gene *tmkB* mutant of *T. virens* IMI 304061 that was also constitutive conidiation in dark as well as in liquid shake culture. Based on the above results, we speculate that the light sensing pathway might be repressed in *Trichoderma* by deletion of Slt2-MAPK genes, but this needs more detailed data to confirm. Slt2-MAPK regulated cell-wall integrity were verified in multiple fungi, including of *Trichoderma* spp. [29, 38], *Metarhizium robertsii* [39], *Penicillium digitatum* [40], and *Saccharomyces cerevisiae* [27]. However, two types of MAPK (Fus3 and Slt2) of strain TC967 were both involved in CWI. The Slt2- and Hog1-MAPK involved in regulating radial growth, the growth became noticeably slower than the wild type after deletion the two types of MAPK genes. The function of Slt2- and Hog1-MAPK in biocontrol acquires few attention because of these gene mutants having poor growth, it was deemed to preclude successful antagonism to pathogens [10]. It is by no means that slow growth rate leads to lose biocontrol ability. Kumar *et al.* first discovered that the Slt2-MAPK gene mutants of *T. virens* strain IMI 304061 retained its ability to overgrow *R. solani* and *Pythium* spp., like the phenotype exhibited by the Fus3-MAPK gene mutants [29]. Our data also showed that the Slt2-MAPK gene mutants had a good biocontrol ability in pot experiment even though it had attenuated growth (Fig 8). The Hog1-MAPK involved in controlling hyperosmotic stress response, our observations exhibited that the growth of Hog1-MAPK gene deletion mutants was totally inhibited on PDA medium added with NaCl, suggesting that *tbmk3* is an essential gene for participating in high osmolarity resistance.

Mycoparasites produce cell wall-degrading enzymes which hydrolyze cell walls of other fungi serving as nutrients for their own growth [9]. We investigated the mycoparasitic strain TC967 and measured the FCWDEs activities of the three MAPK gene mutants. The Slt2- and Hog1-MAPK gene mutants had attenuated ability to overgrow the pathogen *R. solani*, and especially the latter which almost lost antagonism, while the Fus3-MAPK gene mutants retained the ability to overgrow the pathogen. Our findings are in-line with the study on the *tmkB* mutants of *T. virens* IMI 304061 which had relatively weak ability against *S. rolfsii* comparing with the *tmkA* mutants [29]. When the activities of chitinase, protease and β-1,3-glucanase of strain TC967 were measured, that of chitinase and protease were significantly elevated in the Fus3- and Slt2-MAPK gene mutants, which demonstrates that they act as negative regulators. Similarly, the FCWDEs activity enhanced by deletion of Fus3-MAPK gene in *T. virens* Gv29-8 [21] and *T. asperellum* T4 [41]. Except for Fus3-MAPK negatively regulates the related gene expressions of cell wall-degrading enzymes [41], we guess that more FCWDEs have been secreted by the cells of *Trichoderma* strains, because CWI was broken by deletion of Fus3-MAPK gene.

A large amount of VOCs emitted from the *Trichoderma* strains were detected, including simple hydrocarbons, heterocycles, aldehydes, ketones, alcohols, phenols, thioalcohols, thioesters, and their derivatives [42]. The VOCs play a role in communication and inhibition of growth of the target fungi [43, 44]. The previous studies have showed that the VOCs detrimental to plant pathogens *Botrytis cinerea* [45], *Fusarium oxysporum* [46], *R. solani*, *S. rolfsii, Sclerotinia sclerotiorum* [47], and *Alternaria alternata* [48]. In our work, the VOCs emitted from the wild strain TC967 obviously inhibited the mycelial growth of *R. solani*. The inhibitory effect of producing VOCs was further enhanced by deletion of Fus3-MAPK gene. It is possible that the type or composition of effective VOCs for biocontrol changed upon deletion of the MAPK genes. To our knowledge, the interaction of MAPK with VOCs in *Trichoderma* has not been reported. Further intensive researches are surely needed to explore the functions of MAPK involved in regulating VOC syntheses.

The intercommunities and divergences in regulation of biocontrol in one strain mediated by three different types of MAPK could be attributed to broad factors. The expressions of downstream genes that are regulated by these MAPKs ultimately determine the phenotypes of fungi. However, the regulatory process is very complex, the cross-talk interaction of the two MAPK pathways were found in some fungi [40, 49], even the MAPK pathways united other signal pathways co-regulate downstream gene expressions in *Metarhizium* species [50]. Using transcriptomic analysis at the genome-wide level is a valuable tool to identify downstream genes that are regulated by signaling pathways. RNA-seq has been used to reveal the gene expression changes regulated by Fus3 and Slt2-MAPK in *T. reesei* during sugarcane bagasse degradation [51], but using RNA-seq to explore *Trichoderma* whole gene expression regulated by MAPK in biocontrol is lacking. In the present study, we predicted the composition of biocontrol-related genes and systematically investigate the gene expressions at genomic level that are regulated by Fus3-, Slt2- and Hog1-MAPK induced by *R. solani* using RNA-seq. The data indicated that the expressions of a large number of genes were changed by deletion of each MAPK gene (Fig 9), which further tells that MAPKs play major roles in regulation of the biocontrol-related genes. In comparison, more genes were upregulated in Fus3-MAPK gene mutants. Differing from Fus3-MAPK gene mutants, Slt2 and Hog1-MAPK gene mutants had a similar number of upregulated and downregulated genes induced by *R. solani*. This means Fus3-MAPK primarily play negative regulatory roles in *Trichoderma* antagonistic against the pathogen.

*Trichoderma* strains are a rich source of secondary metabolites. Mycoparasitic *Trichoderma* strains are enriched in secondary metabolism-related genes compared with non-mycoparasitic strains. NRPS clusters were most diverse and functionally important for fungal secondary metabolites [52]. In this study, 11 NRPS clusters were found in the genome of strain TC967, which is less than those of the mycoparasitic strains of *T. virens* and *T. atroviride* (28 and 16, respectively) but nearly equivalent to the non-mycoparasitic *T. reesei* (10) [52]. The secondary metabolites produced by the wild type TC967 did not show obvious inhibition effects on *R. solani* (Fig 6 A and B). However, inhibition ability was promoted by the deletion of Slt2-MAPK gene (Fig 6 A and B). The possible reason is that it changes the content of secondary metabolites. It was also proved that most of the core genes related to secondary metabolism were upregulated in Slt2-MAPK gene mutants (Fig 10A).

FCWDEs are important for the survival of fungi in nature [53]. As a mycoparasitic fungus, *T. brevicrassum* strain TC967 should have a powerful system of FCWDEs to attack *R. solani*. The number of FCWDE genes in strain TC967 (88) is similar to that of the other mycoparasites in the same genus, *T. atroviride* (89), *T. harzianum* (97), and *T. virens* (97) [53]. The GH18 family (chitinases) is considered to be critical and essential during mycoparasitism [54, 55]. The largest number of FCWDEs was GH18 family in the genome of strain TC967 (S3 Table), this means strain TC967 has strong chitinase activity. In most non-mycoparasitic ascomycetes, such as *S. sclerotiorum*, *B. cinerea*, *B. graminis*, *Phaeosphaeria nodorum*, *Magnaporthe oryzae*, *Neurospora crassa* and *Aspergillus niger*, the GH16 family (endo-β-1,3-glucanase) was the most abundant in FCWDEs [56]. However, the wild strain TC967 showed low chitinase activity, the Fus3- and Slt2-MAPK gene mutants appeared to have higher chitinase activity (Fig 7). The result of RNA-seq indicated that expression of the chitinase gene EVM0010536 (*ech42*-like) was obviously upregulated, and expression of GH16 family gene was downregulated in each of the three MAPK gene mutants (Fig 10B). This again says that MAPKs play an important role in regulating expression of FCWDE-related genes.

As a group of small secreted cysteine-rich proteins, the proportion of SSCPs in all proteins of strain TC967 was 7.7% that was similar to some other fungi, such as 5.5% in *Sclerotinia* spp., 6.5% in *Botrytis* spp., 6.7% in *Trichoderma* spp., and 9.4% in *Paraphaeosphaeria sporulosa* [53]. Functions of SSCPs were mostly focused on fungus-plant interactions in the previous researches [15, 57–59]. The function of SSCPs in biocontrol lacks understanding. We analyzed the expressions of SSCP genes in the MAPK gene mutants induced by *R. solani*, most of them were upregulated in Hog1-MAPK gene mutants. And therefore, MAPK involved in regulating expressions of SSCP genes. Further investigation is necessary to confirm the functions of SSCPs in interaction between mycoparasitic *Trichoderma* and phytopathogen.

## Materials and methods

### Fungal strains

*Trichoderma brevicrassum* strain TC967 was isolated from soil of the Tibet Autonomous Region. The phytopathogen *Rhizoctonia solani* (ACCC 36124) was by the courtesy of Professor Yong-Chun Niu, which was isolated from diseased root of spinach. The wild type strain, resultant gene deletion and complemented strains of strain TC967 and *R. solani* were cultured on potato dextrose agar (PDA) medium (200 g potato, 20 g dextrose, 20 g agar, and 1 L water) at 25°C unless otherwise indicated.

### Genomic sequencing and analysis

Genomic DNA (gDNA) of strain TC967 was extracted from 4-day-old mycelia following the protocol described in the handbook of the Plant Genomic DNA Kit (Tiangen, Beijing, China). The gDNA was sequenced by Illumina HisSeq 2500 sequencing (Illumina, CA, USA). The sequence reads were assembled using Velvet [60] and SSPACE [61]. *Ab initio* gene prediction was performed using Genscan [62], Augustus (version 2.4) [63], GlimmerHMM (version 3.0.4) [64], GeneID (version 1.4) [65] and SNAP (version 2006-07-28) [66]. Using GeMoMa (version 1.3.1) [67] predicted genes based homologous protein. Combining two prediction results as mentioned above were performed using EVM (version 1.1.1) [68].

### Retrieval and phylogenetic analysis of *mapk* sequences

The sequences of TMK1 (XP_013957486), TMK2 (XP_013959852), and TMK3 (EHK21342) of *T. virens* Gv29-8 were downloaded from protein database of NCBI (https://www.ncbi.nlm.nih.gov/protein). MAPK genes of strain TC967 were performed against the TMK1, TMK2 and TMK3 proteins by BLASTP (e-value < 1 × 10^-10^, identity > 70%).

ClustalX was used to generate multiple sequence alignments. Phylogenetic analysis was performed based on full-length protein sequences using MEGA-X program by the maximum-likelihood (ML) method with a bootstrap value of 1000 replications.

### Construction of *mapk* gene deletion and complementation mutants

The *mapk* gene mutants based on homologous recombination mediated by *Agrobacterium tumefaciens* AGL-1 was conducted as the previously described [7]. The primers used in construction mutants were listed in S5 Table. For each *mapk* gene, two pairs of primers were designed to amplify approximately 2 kb of the 5′ and 3′ flanking sequences. The hygromycin-resistance gene (*hygR*) cassette in pCAM-eGFP plasmid [7] was amplified by PCR. The *hygR* cassette was inserted into 5′ and 3′ flanking sequences of each *mapk* gene subsequently. The *hygR* cassette and flanking sequences were ligated with the linearized pCAMBIA2301 plasmid to produce gene mutant plasmid pCAM-flank-*Δtbmk1*, pCAM-flank-*Δtbmk2* and pCAM-flank-*Δtbmk3*. To complement gene disruption mutants, the *hygR* cassette of each gene mutant plasmid was replaced by *mapk* linked neomycin resistant gene (*neoR*) to construct the gene complementation plasmid. All the fragments were ligated using One Step Cloning Kit (Vazyme Biotech, China). The recombinational plasmid were verified by sequencing to ensure the accuracy of the in-frame fusion region. The colonies of *mapk* deletion mutants were observed on PDA plates supplemented with 120 µg/mL hygromycin B. The colonies of *mapk* complement mutants were screened on PDA plates supplemented with 1200 µg/mL neomycin sulphate. The resulting mutants were confirmed by PCR assays, primers were listed in S5 Table.

Southern blotting was used to detect the *hygR* cassette copy number in positive mutants. gDNA of *Δtbmk1*, *Δtbmk2* and *Δtbmk3* were digested by *Bst*XI, *Bfu*I and *Bst*XI, respectively (S2 Fig). Products were separated by 1.1% agarose gel electrophoresis and hybridized with Biotin-labeled DNA probe. DNA fragments as probe1, probe2 and probe3 were amplified from gDNA of TC967 using primer sets sou1-F/R, sou2-F/R and sou3-F/R (S5 Table), respectively. A LumiPRO ECL substrate kit (Cleaver Scientific, United Kingdom) was used for visualization of nucleic acid bands.

### Growth and conidiation

To assess differences in growth among strains, size and morphology of the colonies were assessed daily. For recording conidial yields, the strains were incubated in 90 mm PDA Petri dishes at 25°C under a light/dark cycle or only in dark condition for 7 days. The conidiospores were washed by 5 mL distilled water and filtrated through two layers of Miracloth (Calbiochem, Germany). The yield of spores was measured by a hemocytometer. Hyphae and conidiophore of the strains were observed under optical microscope (Zeiss Imager A2, Germany). Each experiment was repeated three times.

One milliliter conidial suspension (1.0 × 10^5^ spores per mL) of the strains were added into 250 mL flasks containing 100 mL potato dextrose broth (PDB) and inoculated for 48 h at 25°C with 150 rpm shake in dark condition. The experiment was repeated three times.

### Tolerance to abiotic stresses

Five mm diameter fresh mycelium plugs of the *Trichoderma* strains were inoculated on PDA plates containing 300 mg/L Congo red (CR), 0.5 M NaCl, and 30% H_2_O_2_, respectively. These plates were grown at 25°C for 3 days prior to examination. Colony diameters of individual samples were measured for comparison. Three replicates of each experiment were set up.

### Dual cultures

The dual culture method followed our previously study [7]. A 5 mm diameter mycelium plug of *Trichoderma* was put on PDA plates with that of *R. solani* 5 cm apart on the same plates. Plates only inoculated with mycelium plug of *R. solani* served as the control. Three replicates were set up for each treatment, and all plates were incubated at 25°C with a 12 h light/dark cycle for 7 days.

### Growth inhibition by non-volatile secondary metabolites

Non-volatile secondary metabolites of *Trichoderma* wild and mutant strains were tested in plate assays [69] with some modifications. A 5 mm diameter mycelium plug of the *Trichoderma* strains was inoculated on 9 cm PDA plates covered with a sterile cellophane disc. The cellophane disc was removed when *Trichoderma* colony covered about 3/4 of the plate, and then a 5 mm diameter mycelium plug of *R. solani* was placed in the middle of the plate. Colony diameter of *R. solani* was recorded after 48 h, and the growth inhibition was calculated by comparing with that of *R. solani* on PDA. Data was generated from three replicates of each experiment.

### Extracellular hydrolase activity

The method of measuring chitinase activity of the strains was as our previously described [7]. For determination of protease activity, a mycelium plug of individual *Trichoderma* strains was inoculated on the skim milk agar medium (skim milk 50 g/L, agar 15 g/L) and incubated for 2 days at 25°C, then proteolytic circle was observed.

The 1 mL spore suspension (1.0 × 10^7^ spores per mL) of each *Trichoderma* strain was cultured in TLE medium [bactopeptone 1.0 g/L, urea 0.3 g/L, KH_2_PO_4_ 2.0 g/L, (NH_4_)_2_SO_4_ 1.4 g/L, MgSO_4_·7H_2_O 0.3 g/L, CaCl_2_·6H_2_O 0.3 g/L, trace elements (Fe^2+^, Mn^2+^, Zn^2+^, and Co^2+^) 0.1%, glucose 0.2 g/L, cell walls powder of *R. solani* 5 g/L] [70] in a rotary shaker at 25°C and 150 rpm for 7 days. After 48 h, the culture filtrate was collected every day by centrifuging and used as a source of enzymes. Chitinase activity were determined using the chitinase activity assay kit (Boxbio, China), 1 U of chitinase activity was defined as the amount of enzyme required to release 1 μmol N-acetylglucosamine per hour. Protease activity was measured by the azocasein procedure following the reference [71], one enzyme unit was defined as the amount of enzyme required to elevate one unit of absorbance per hour. β-1,3-glucanase activity was determined using laminarin as a substrate, the detailed protocol following the reference [72]. One enzyme unit was defined as the amount of enzyme necessary to release 1 mg reducing sugar per 30 minutes.

### Growth inhibition by volatile organic compounds (VOCs)

The *Trichoderma* strain and that of *R. solani* were inoculated in a 9 cm Petri dish divided into two compartments (I-plate). Each of the two compartments of the I-plates was filled with ca. 8 mL PDA. A 5 mm diameter mycelium plug of a *Trichoderma* strain was inoculated on one side of a I-plate. Two days later, the other side of the I-plate was inoculated with a 5 mm diameter plug of *R. solani*. In the control treatment, *R. solani* mycelium plug was inoculated in one compartment of the I-plate with the absence of *Trichoderma*. The I-plates prevent the contact between *Trichoderma* and *R. solani*, but allow free exchange of VOCs. The I-plates were sealed with two layers of Parafilm to prevent from VOCs leaking, and incubated at 25°C for 48 h. There are four replicates for each treatment. Mycelial growth inhibition of *R. solani* was measured as a differential percentage compared with the control.

### Pot experiments

Surface sterilized cucumber seeds were pregerminated in 150 mm Petri dishes at 25°C. Six treatments were set as follows: Control, not inoculated with any fungi; Rs, medium only inoculated with 1.5 g biomass of *R. solani* per kilogram; *Trichoderma* + Rs, medium respectively pretreated with the wild and mutant strains (including TC967, *Δtbmk1*, *Δtbmk2* and *Δtbmk3*) and then inoculated with *R. solani*. Conidia of *Trichoderma* (1.0 × 10^6^ per gram medium) were previously inoculated in plastic pots (8 cm length × 8 cm width × 11.5 cm high) with about 200 g sterilized podsolic soil and vermiculite 1:1 (v/v). One week later, the cucumber plants were planted into the pots. Three weeks after transplanting, the treatments of Rs and *Trichoderma* + Rs were inoculated with 15 mL *R. solani* mycelium homogenated around cucumber stem base. All pots were kept in greenhouse at temperature of 25/20°C with 16/8 h day/night photoperiod, and maintained 70% relative humidity using humidifier. Two weeks later, disease symptoms were scored as 5 classes (0−4): 0, healthy; 1, 1−25% of stem rot; 2, 26−50% of stem rot; 3, 21−75% of stem rot; and 4, more than 75% of stem rot or plant dead. Then, a disease index (DI) was calculated by the formula: DI = [(1n_1_ + 2n_2_ + 3n_3_ + 4n_4_)/4N] × 100%, n_1_−n_4_ is the number of plants with score 1−4, respectively, and N represents the total number of plants used in the variant [73].

### Gene clusters and gene identifications

The antiSMASH (fungal version) with default parameters was used to predict the secondary metabolite biosynthesis gene clusters of strain TC967 [74].

The dbCAN server was conducted against the sequence libraries and profiles in the carbohydrate-active enzymes (CAZy) database. The matched hits were chosen using HMMER program with e-value < 1e^-15^ and coverage > 0.35. Fungal cell wall-degrading enzymes (FCWDEs) were classified based on the active domain in the CAZy database following Zhao et al. [75].

The secretory signal peptide of strains TC967 were searched using SignalP 5.0 services (https://services.healthtech.dtu.dk/service.php?SignalP-5.0) with default parameters for eukarya. The small secreted cysteine-rich proteins (SSCPs) were chosen as less than 300 amino acids and more than 4% cysteine content [53].

### Transcriptomic analysis

The *Trichoderma* strains were grown on PDA plates covered with cellophane at 25°C and 12 h cyclic illumination. The mycelia were harvested after colony of *Trichoderma* overgrowing that of *R. solani* by ca. 5 mm. Three replicates for each treatment were performed. The mycelial samples were frozen in liquid nitrogen immediately. Total RNA was extracted from the mycelia using RNeasy Plant Mini Kit (Qiagen, Germany). The quality of the extracted RNA was assessed on 0.9% agarose gel. The RNA purity, integrity and concentration were evaluated using Nanodrop 2000 (Thermo Fisher Scientific, Massachusetts, USA) and Agilent 2100 (Agilent Technologies, State of California, USA). Each RNA sample was sequenced on an Illumina NovaSeq 6000 (Illumina Inc., San Diego, CA, USA) with 150 bp paired-end reads at Biomarker Co. Ltd (Beijing, China).

Transcripts were generated by mapping the RNA-seq reads to reference genome using Hisat2 with default parameters [76]. Differential expression analysis was performed using StringTie with default parameters [77]. DESeq2 (v1.6.3) was used to analyze the differentially expressed genes (DEGs) [78]. Fragments per kilobase of transcript per million fragments mapped (FPKM) values were served as measurement units to estimate the expression level of each gene. A false discovery rate (FDR) < 0.01 and |log_2_(Fold Change)| ≥ 1.5 were set as the thresholds for DEGs identification. The Venn diagram drawing and heatmap were executed by TBtools [79]. The GO enrichment analyses of DEGs were carried out using BMKCloud (http://www.biocloud.net/).

### Real-time qPCR analysis

Total RNAs (1000 ng per reaction) were transcribed into complementary DNAs (cDNAs) with HiScript^®^ III RT SuperMix for qPCR (+gDNA wiper) (Vazyme Biotech, China). The real-time qPCR was conducted in 20 μL reaction mixtures containing 1.0 μL 10-fold dilution cDNA, 0.8 μL each of the forward and reverse primers (10 μM), 10 μL 2 × ChamQ Universal SYBR qPCR Master Mix (Vazyme Biotech, China), and processed using CFX96 real-time qPCR detection system (Bio-Rad, USA) with the following procedure: denaturation at 95°C for 30 s followed by 40 cycles of denaturation at 95 °C for 10 s, and extension at 60 °C for 30 s. Primers used in real-time qPCR analysis were presented in S6 Table. The relative normalized transcript level of each gene was computed using the 2^−ΔΔCt^ method [80]. Data for the three biological replicates were assayed in triplicate for each gene.

## Author contributions

YZ and W-YZ conceived and designed the experiments. ZY performed the experiments and wrote the manuscript. W-YZ revised and finalized the manuscript.

## Competing interests

The authors declare no competing interests.

## Funding

This project was supported by the National Natural Science Foundation of China (31570018) and National Key Research and Development Program (2017YFD0200600).

## Acknowledgements

We are grateful to Dr. Kai Chen for isolating *Trichoderma brevicrassum* strain TC967, Prof. Yong-Chun Niu for providing strain of *Rhizoctonia solani*, Prof. Hui-Shan Guo and Dr. Hua-wei Wu for laboratory facilities and technique assistance, and Prof. Hao Lu for providing vectors of construction of *Trichoderma* mutants.

## Supporting information

**S1 Fig. Multiple alignments and domain analyses of MAPK proteins of *Trichoderma brevicrassum* strain TC967.** Highlighted part shows the conserved signature motif obtained with Clustal X program. The red boxes indicate the conserved motif in three types of MAPK. GenBank accession numbers were as the follows: *Blumeria graminis* MAP1 (AAG53654) and MAP2 (AAG53655); *Botrytis cinerea* BMP1 (AAG23132) and BcSAK1 (AM236311); *Claviceps purpurea* CPMK1 (CAC47939) and CPMK2 (CAC87145); *Saccharomyces cerevisiae* FUS3 (AAA34613), SLT2(CAA41954) and HOG1(AAA34680); *T. atroviride* TMK1 (XP_013941981) and TMK3 (XP_013941593); *T. brevicrassum* strain TC967 TbMK1 (EVM0011518), TbMK2 (EVM0003677) and TbMK3 (EVM0007810); *T. guizhouense* SPM1 (OPB43978); *T. reesei* TMK1 (XP_006965066), TMK2 (XP_006969672) and TMK3 (XP_006962041); *T. virens* TMK1 (XP_013957486), TMK2 (XP_013959852) and TMK3 (EHK21342).

(TIF)

**S2 Fig. Disruption of *mapk* genes in *Trichodrema brevicrassum* strain TC967.** (A) Schematic representation of deleting *tbmk1*, *tbmk2* and *tbmk3* genes. (B) PCR assays for verification of wild and gene deletion mutant strains. In each *mapk* gene, three randomly independent disruption mutants were chosen. (C) Southern blotting analysis of transformants. gDNA of each *mapk* mutants were digested by restriction endonuclease showed in (A). The blots were hybridized with the probes indicated in (A). (D) PCR to verify the complementation mutants. C-1 and C-2 were two randomly independent complemented mutants. The primers of construction deletion plasmids and confirmation of mutants were shown in S5 Table.

(TIF)

**S3 Fig. Dual cultures of *Trichoderma brevicrassum* strain TC967 *mapk* mutants against *Rhizoctonia solani*.** The plates were incubated in a 25°C incubator with cycle light for 7 days.

(TIF)

**S4 Fig. CWEDs activity of *Trichoderma brevicrassum* strain TC967 *mapk* mutants in liquid fermentation during 2−7d.** (A) Chitinase, (B) Protease and (C) β-1,3-glucanase. The bars represent the averages of standard deviations of three biological replicates.

(TIF)

**S5 Fig. Heatmap of correlation among expressions of *Trichoderma brevicrassum* strain TC967 samples.** Spearman’s correlation coefficient R was applied to evaluate reproducibility of biological replicates. A closer R^2^ value to 1 indicates better reproducibility between the two samples.

(TIF)

**S6 Fig. Validation of expression of randomly selected DEGs using RNA-seq and real-time qPCR analyses for *Trichoderma brevicrassum* strain TC967 mutants *Δtbmk1* (A), *Δtbmk2* (B) and *Δtbmk3* (C) induced by *R. solani,* respectively.** The log_2_(Fold Change) is calculated from the mean of three replicates. The coefficient of determination (R^2^) is displayed. The randomly selected DEGs included in nonribosomal peptide synthetase (EVM0000012), glutathione-dependent formaldehyde-activating enzyme (EVM0001330, EVM0001393, EVM0011384), BrlA (EVM0001742), function unknown (EVM0001929), chitin synthase (EVM0002032), glycosyl hydrolase (EVM0002519), hydrophobin (EVM0002575), β-1,3-glucan synthase (EVM0003016), β-ketoacyl synthase (EVM0004965), cyclochlorotine synthetase (EVM0005196), RDR1 (EVM0005281), FluG (EVM0007122), chitin deacetylase (EVM0008955), glutamine synthetase (EVM0010151), chitinase (EVM0010536), trihydrophobin (EVM0011084), glucanase (EVM0011451), and β-ketoacyl synthase gene (EVM0011588).

(TIF)

**S7 Fig. Core biosynthetic genes related to secondary metabolism of *Trichoderma brevicrassum* strain TC967.** (A) Structures of core NRPS, NRPS-like and T1PKS-NRPS biosynthetic proteins; (B) Structures of core T1PKS biosynthetic proteins. C, condensation domain; A, adenylation (AMP) domain; PP, peptidyl-carrier protein domain; TD, thioesterase domain; KR, ketoreductase; NAD, nicotinamide adenine dinucleotide; ER, enoyl reductase domain; DH, polyketide synthase dehydrogenase; cMT, cycle methyltransferase; SAT, serine acetyltransferase; TE, thioesterase.

(TIF)

**S1 Table Comparisons of the sampled *Trichoderma brevicrassum* strain TC967 sequencing data with reference genome sequence**

(XLSX)

**S2 Table Distribution of cluster of secondary metabolism in *Trichoderma brevicrassum* strain TC967**

(XLSX)

**S3 Table Composition of fungal cell wall-degrading enzymes in *Trichoderma brevicrassum* strain TC967**

(XLSX)

**S4 Table GO enrichment analysis of SSPs in *Trichoderma brevicrassum*strain TC967**

(XLSX)

**S5 Table The primers of constructing *mapk* mutants**

(XLSX)

**S6 Table The primers of Real-time qPCR**

(XLSX)

## Notes

### Competing Interest Statement

The authors have declared no competing interest.

